# Harnessing TfR1 for Cross-Species Systemic Delivery of siRNAs to Deep Brain Regions Using Single-Domain Antibodies

**DOI:** 10.64898/2026.05.20.726486

**Authors:** Guillaume Jacquot, Marion David, Bélinda Pecqueux, Yasmine Mechioukhi, Stéphane D. Girard, Magali Godard, Karine Varini, Cléa Boursery, Cécile Frapolli, Sébastien Roux, Martin Bigonnet, Béatrice Brousse, Emma Augustin, Gersande Godefroy, Christophe Fraisier, Bastien Serrano, Andréa Romette, Morgane Thomas, Kader Mazouzi, Baptiste Calleya, Diane Beuzelin, Aude Faucon, Karima Abouzid, Gauthier Dangla Pélissier, Pascaline Lécorché, Soioulata Aboudou, Florian Benoit, Maxime Masse, Géraldine Ferracci, Jamal Temsamani, Michel Khrestchatisky

## Abstract

Despite their therapeutic potential across a wide range of central nervous system (CNS) disorders, nucleic acid-based therapeutics are limited by inefficient delivery to deep brain regions at clinically viable doses. Transferrin receptor 1 (TfR1) has emerged as an attractive target for receptor-mediated transcytosis across the blood–brain barrier (BBB), enabling systemic delivery of biologics such as lysosomal enzymes and monoclonal antibodies. In this study, we demonstrated the translational potential of recently described TfR1-targeting camelid-derived single-domain antibodies (VHHs) for CNS delivery of siRNAs. When conjugated 1:1 to different tool siRNAs, these VHHs promote rapid and robust intracellular uptake, resulting in potent RNAi activity at low nanomolar concentrations in neural cells. Systemic administration of VHH-siRNA conjugates in wild-type mice, hTfR1 transgenic-mice and non-human primates revealed a favourable pharmacokinetic profile characterized by rapid TfR-dependent distributional clearance and efficient functional uptake in deep brain structures. This translated into durable target knockdown of 50–80% at both mRNA and protein levels and with ED50 below 1 mg/kg siRNA. Collectively, these findings establish our TfR1 targeting VHHs as a fit-for-purpose platform for the systemic delivery of therapeutic oligonucleotides to deep brain structures at clinically relevant doses, opening new avenues for the treatment of diverse CNS disorders.

**Graphical abstract:** 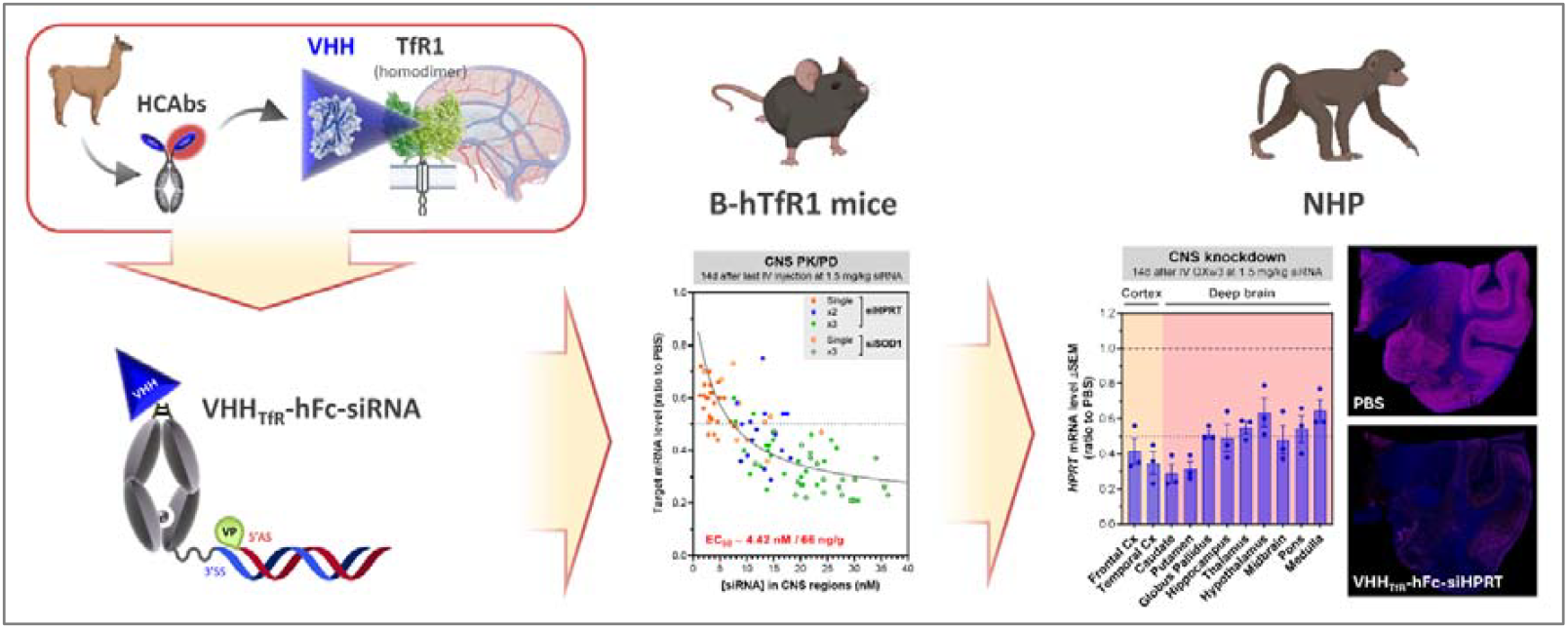

## Introduction

Over the past two decades, genetic and precision medicine have attracted considerable interest as important therapeutic strategies for the prevention and treatment of disease by addressing underlying molecular mechanisms. In particular, modulation of gene expression using oligonucleotide therapeutics (OTs), including antisense oligonucleotides (ASOs) and small interfering RNAs (siRNAs), have progressed from experimental tools to approved medicines. Since the first approval of fomivirsen in 1998, more than 20 RNA-based drugs have reached the market with hundreds of additional candidates currently in clinical development, including approximately 10 that target central nervous system (CNS) indications (1-6). Huge efforts in the optimization of OTs design and chemistry were key to this achievement (6-8). Unlike conventional drugs that mostly target proteins with a one-size-fits-all approach, OTs offer a means to tackle the underlying genetic causes of diseases and offer critical advantages: they achieve high target specificity thanks to Watson-Crick base pairing, and prolonged pharmacodynamic effects owing to their high metabolic stability, allowing infrequent dosing. Finally, OTs also have the potential to fill the gap between rather common disorders to ultrarare and even patient-specific genetic conditions, as illustrated by the single patient-customized milasen (9,10).

Despite their huge therapeutic potential, the clinical translation of OTs remains limited due to their poor pharmacokinetics (PK), low and non-specific tissue distribution and restricted access to intracellular sites of action (11-15). To date, the most efficient and inspiring technology consists in conjugating OTs to the triantennary N-acetylgalactosamine (GalNAc), enabling efficient and specific uptake in liver hepatocytes *via* the asialoglycoprotein receptor (ASGPR) (16,17). This seminal work rapidly translated into clinical success in liver diseases (18), and has spurred the identification of new receptor-ligand pairs with similar functional delivery potential in extrahepatic tissues (11). The most challenging target organ and disease area for OTs is the CNS, comprising the brain, spinal cord and retina. These vital organs are protected from potentially harmful compounds and pathogens by specialized barriers, including the blood–brain barrier (BBB) and blood–retina barrier (BRB) (19-21). However, these same barriers significantly hinder OTs from reaching the nervous tissue parenchyma.

As a result, CNS-directed OTs in development pipelines rely mostly on local administration routes such as intrathecal (IT) delivery into the cerebrospinal fluid (CSF) for brain and spinal cord indications, or intravitreal (IVT) injection for retinal diseases (7). While these approaches have enabled clinical efficacy, they present several limitations, including high concentration gradients between the injection site and distant regions, requiring high and frequent dosing to achieve meaningful pharmacodynamic response in deep brain structures (22,23). In addition, both IT and IVT delivery are highly invasive and associated with procedure-related adverse effects, which may limit long-term tolerability and patient compliance (7,21,24,25). Moreover, effective tissue uptake following local administration relies largely on small molecular size and high phosphorothioate (PS)-content, which promote non-specific hydrophobic interactions with cell-surface proteins, and uptake *via* a process called ‘gymnosis’ (26-28). This dependence restricts the range of OT modalities that can be effectively delivered, particularly double stranded siRNAs or constructs with reduced PS content. Although alternative strategies aimed at enhancing tissue retention, such as lipid conjugation or divalent siRNA architectures, have shown promise (5,29), receptor-mediated uptake offers a potentially more generalizable solution.

Beyond acting as a safeguard interface, the BBB regulates brain homeostasis by selectively transporting essential nutriments *via* receptor-mediated mechanisms expressed at the luminal (blood) side of brain endothelial cells (BECs). Due to the high vascular density of the CNS, most neural cells are located within approximately 20 µm from a capillary (30), making systemic delivery via receptor-mediated transcytosis (RMT) an attractive strategy to achieve widespread CNS distribution (31). Among BBB-expressed receptors, transferrin receptor 1 (TfR1) is one of the most extensively characterized and is involved in CNS supply of transferrin-bound iron (32-34). Therapeutic exploitation of TfR1-mediated RMT has recently reached the clinic, as demonstrated by the approval of a TfR1-targeting bispecific protein for enzyme replacement therapy in Hurler’s Syndrome, in Japan (35). The first preclinical proof-of-concept that this receptor is also able to mediate oligonucleotide delivery *in vivo* came from Sugo and colleagues (36). In this seminal work, the authors demonstrated that an anti-TfR1 Fab’-siRNA conjugate, targeting the hypoxanthine-guanine phosphoribosyltransferase (HPRT), induced durable knockdown in muscle following intravenous administration in mice. Since then, several TfR1-targeting antibody- or Fab-based ligand-OT conjugates, collectively referred to as Antibody-Oligonucleotide Conjugates (AOCs), have entered early clinical development, primarily for muscle indications (1,37). Recently, full-length IgG-based technologies (□150 kDa) have demonstrated that the BBB-expressed TfR1 can mediate the delivery of multiple classes of OTs to the CNS in preclinical models (38). These approaches include: (i) a splicing-modulating PMO (phosphorodiamidate morpholino oligonucleotide) targeting the *SMN2* (survival motor neuron 2) primary transcript, delivered using the mouse TfR1-binding 8D3 antibody variant (8D3_130_) (39); (ii) a full-PS gapmer antisense oligonucleotide (ASO) targeting the long non-coding *MALAT1* mRNA (metastasis-associated lung adenocarcinoma transcript 1), conjugated to a human/non-human primate (NHP) cross-reactive antibody in the Oligonucleotide Transport Vehicle (OTV) format (40); and (iii) siRNAs targeting *SNCA* (α-synuclein) or *MAPT* mRNAs, also delivered using a human/NHP cross-reactive antibody (41). Additionally, promising results have been reported with a reduced-size antibody-based Brainshuttle, site-specifically conjugated via bacterial transglutaminase to an ASO targeting *MAPT* mRNA (microtubule-associated protein tau) (42). Nevertheless, further reductions in conjugate scaffold size, coupled with enhanced CNS delivery efficiency, remain necessary to achieve robust pharmacodynamic effects at clinically relevant doses.

We recently identified and optimized a panel of more than 150 single-domain antibody (sdAbs or VHH) variants, derived from initial C5 and B8 hits, that bind TfR1 with cross-reactivity in rodents, NHPs and human (43-46). This diversification effort yielded approximately 30 variants exhibiting less than a 2-fold affinity difference between NHP and human TfR1. These VHHs are derived from camelid heavy chain-only antibodies (HCAbs) and consist of a single variable domain with a molecular weight of *ca*. 12–15 kDa (47). These VHH bind a previously unexploited epitope within the hinge region of the TfR1 homodimer. Owing to their small size, favorable biophysical properties, and ability to access cryptic epitopes, VHHs are increasingly used as targeting ligands in both preclinical and clinical settings (48), and represent a promising AOC approach for targeted delivey of OTs (37,38,49).

In this study, we investigated whether our unique TfR1-targeting VHHs can function as delivery ligands for systemically administered oligonucleotides, enabling transport across the BBB and uptake into CNS parenchymal cells. To avoid confounding effects associated with gymnotic uptake, we focused on the siRNA modality, although single-stranded full-PS ASOs are presently the most common oligonucleotide modality in CNS clinical pipelines (1,6). We selected previously validated siRNAs targeting the ubiquitously expressed *Sod1* gene (superoxide dismutase 1, rodent selective siRNA) (5) and *Hprt* gene (rodent/NHP cross-reactive) (36), allowing unbiased assessment of functional delivery across tissues and species. Using 1:1 defined VHH-siRNA conjugates, we demonstrated efficient TfR1-mediated intracellular delivery *in vitro*, correlating with fast distributional clearance and robust (50-80%) target gene knockdown in deep brain regions following low systemic dose administration in both mice and NHPs.

## Results

### General design and TfR1-binding of VHH-hFc-siRNA conjugates

For initial proof-of-concept studies in rodents and NHPs following systemic administration, we employed a PK-enhanced heterodimeric VHH-hFc-siRNA construct based on the Knob-into-Hole technology (50). This architecture comprises a □14 kDa TfR1-targeting VHH fused at the N-terminus of one crystallizable fragment (Fc) arm of a human IgG1 engineered with LALA-PG mutations (L234A/L235A/P329G) to attenuate FcγR-mediated effector functions (51). The opposing Fc arm was modified at its C-terminus with a Q-tag to enable site-specific conjugation of a single siRNA via bacterial transglutaminase (BTG), followed by copper-free strain-promoted azide-alkyne cycloaddition (SPAAC) click chemistry (**Figure 1**) (52). The resulting conjugates have a molecular weight of *ca*. 81-83 kDa, depending on the VHH variant used and siRNA design, namely 2-fold smaller than conventional AOCs (40,53).

**Figure 1.**
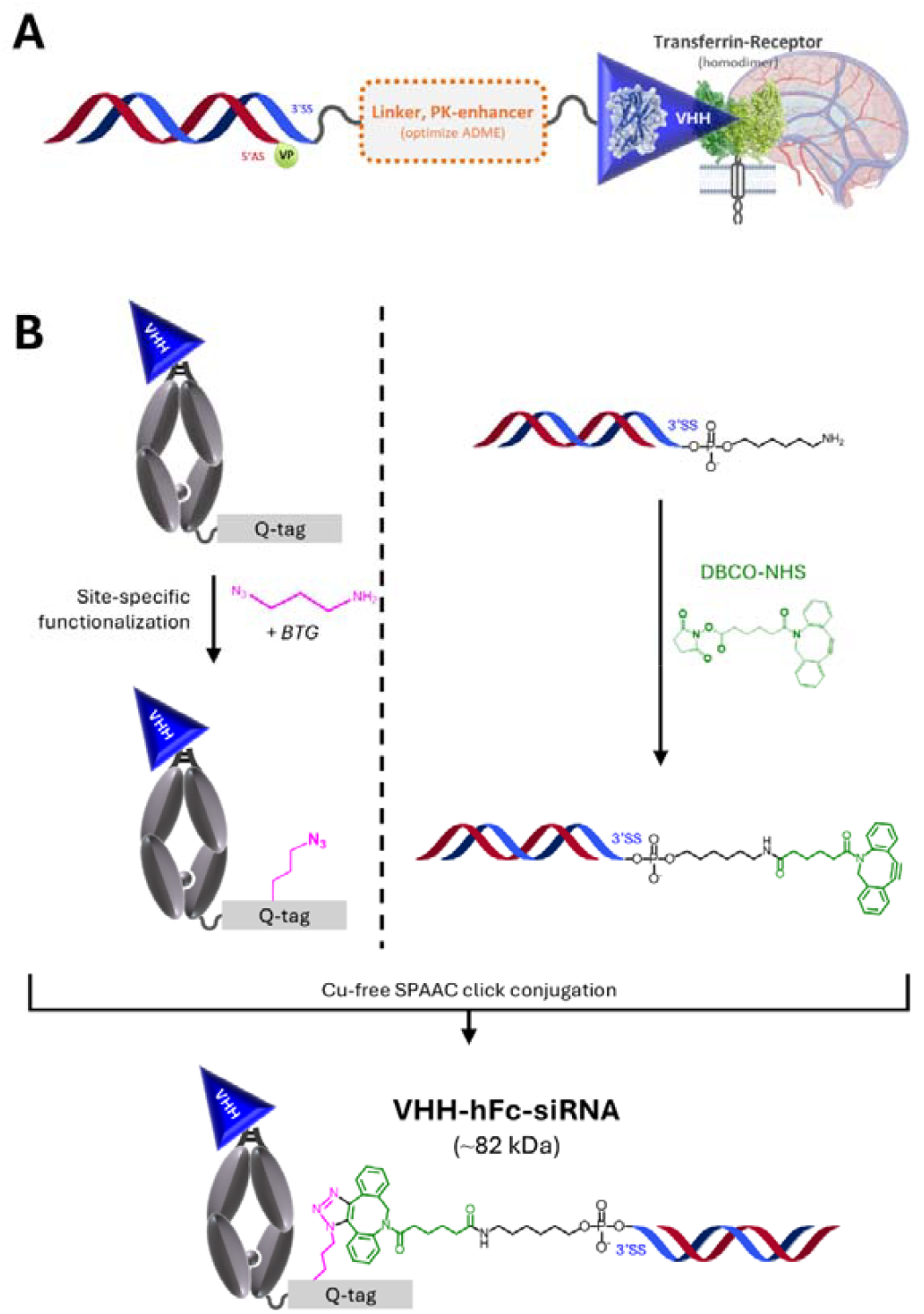
Conjugation strategy used to generate VHH-Oligo and VHH-hFc-Oligo conjugates. (A) Schematic representation of a VHH-Oligonucleotide conjugate design intended for functional delivery in CNS tissues after systemic administration. The present work focuses on siRNAs as they display only low intrinsic uptake potential, allowing unbiased evaluation of our TfR-mediated delivery platform. (B) VHH-hFc-siRNA conjugates used in this work were generated using a two-step approach involving first, functionalization of the siRNA with an alkyne group (exemplified with a DBCO) attached to an aminohexyl group and introduction of an azide group in C-terminus of a heterodimeric VHH-hFc fusion (Fc fragment of a human IgG1 used as a model PK-enhancer) using bacterial transglutaminase; and second, copper-free strain-promoted azide-alkyne cycloaddition (SPAAC) click conjugation. The purified conjugates have a VHH-to-siRNA ratio of 1:1.

From our extensive panel of rodent, NHP, and human cross-reactive VHH variants derived from the C5 and B8 parental hits (44), we selected the C5 and the B8V32 variants for studies performed in wild-type mice (WT) and in hTfR1 ectodomain (ECD)-homozygous mice (B-hTfR1), respectively. Indeed, these variants display similar binding parameters in their free form towards their respective target, with *K*_*D*_ values of 87.4 nM for C5 on mouse TfR1 (mTfR1) and 29.7 nM and 47.6 nM for B8V32 on human TfR1 (hTfR1) and cynomolgus TfR1 (cTfR1), respectively (**Table 1**). As a negative control in both *in vitro* and *in vivo* experiments, we used a non-binding C5 variant (“C5neg”) harboring two point mutations (Y105A and L107A) that result in an almost complete loss of affinity for TfR1 (44). Upon monomeric fusion to the hFc fragment and conjugation to *Sod1*- or *Hprt*-targeting siRNAs (siSOD1 and siHPRT, respectively), a typical 3-to 8-fold lower binding affinity was observed, largely attributed to a slower on-rate based on Surface Plasmon Resonance (SPR). The resulting C5-hFc-siSOD1, B8V32-hFc-siSOD1 and B8V32-hFc-siHPRT conjugates showed comparable binding profiles at pH 7.4 (**Figure 2A**) – with *K*_*D*_ values in the 100–300 nM range for mTfR1 (C5 conjugate) and h/cTfR1 (B8V32 conjugates), and off-rate constants of approximately 0.02 s^-1^ (**Table 1**), allowing unbiased comparison of their PK/PD profile in further *in vivo* studies in mice and NHPs. Such binding parameters were previously shown to correlate with efficient BBB-crossing and brain delivery of these TfR1-targeting VHHs (44), consistent with previous findings that a monovalent binding mode together with moderate binding affinity to TfR1 are key for efficient transcytosis (33,54-56). Importantly, binding at pH 6.0, corresponding to the pH in sorting endosomes, was strongly affected, as exemplified with the □20-fold lower TfR1-binding affinity of the free B8V32 and the B8V32-hFc-siHPRT conjugate, attributed to a □10-fold slower association rate and a slightly faster dissociation rate. Because this pH-dependent reduction in binding to TfR1 was accompanied by 2-fold lower binding responses (data not shown), we hypothesized that a conformational change of the TfR1 epitope could trigger mechanical dissociation of these VHHs.

**Table 1.** TfR1-binding parameters of the naked VHH_TfR_ and VHH-Oligo conjugates used in the present work using either Surface Plasmon Resonance (SPR) on immobilized TfR1 at pH 7.4 (unless specified) or using a competition assay in living murine Neuro-2a or human MCF7 cells.

| Compound name | TfR1 species | Kinetics parameters using SPR | | | $K_{i/app}$ (M) |
| --- | --- | --- | --- | --- | --- |
| | | $k_{on}$ (on-rate)<br>(M<sup>-1</sup> s<sup>-1</sup>) | $k_{off}$ (off-rate)<br>(s<sup>-1</sup>) | $K_D$<br>(M) | |
| C5neg | mTfR1 | - | - | No binding | No binding |
| C5neg-hFc-siSOD1 | mTfR1 | - | - | No binding | No binding |
| C5 | mTfR1 | 1.08E+06 | 4.79E-02 | **8.74E-08** | **3.13E-07** |
| C5-hFc-siSOD1 | mTfR1 | 1.35E+05 | 3.07E-02 | **2.22E-07** | **3.98E-07** |
| C5neg | hTfR1 | - | - | No binding | No binding |
| C5neg-hFc-siSOD1 | hTfR1 | - | - | No binding | No binding |
| C5neg-siHPRT | hTfR1 | - | - | No binding | No binding |
| C5 | hTfR1 | 6.45E+05 | 1.72E-04 | **2.20E-10** | **2.00E-09** |
| C5-siSOD1(A680) | hTfR1 | 1.67E+05 | 1.15E-04 | **6.89E-10** | ND |
| C5-hFc-siSOD1(A680) | hTfR1 | 1.70E+05 | 1.14E-04 | **6.84E-10** | ND |
| B8h1 | hTfR1 | 6.24E+05 | 2.30E-04 | **4.07E-10** | **3.2** |
| B8h1-siHPRT | hTfR1 | 3.31E+05 | 2.38E-04 | **7.20E-10** | ND |
| B8h1-siHPRT(A680) | hTfR1 | 2.55E+05 | 2.07E-04 | **8.13E-10** | ND |
| B8V32 | hTfR1 | 6.92E+05 | 1.82E-02 | **2.97E-08** | **1.45E-07** |
| B8V32 (pH 6.0) | hTfR1 | 7.93E+04 | 2.56E-02 | **4.36E-07** | N/A |
| B8V32-hFc-siSOD1 | hTfR1 | 9.98E+04 | 1.39E-02 | **1.44E-07** | **2.29E-07** |
| B8V32-hFc-siHPRT | hTfR1 | 9.83E+04 | 1.44E-02 | **1.59E-07** | **1.87E-07** |
| B8V32-hFc-siHPRT (pH 6.0) | hTfR1 | 8.74E+03 | 2.25E-02 | **2.70E-06** | N/A |
| B8V32-siHPRT | hTfR1 | 2.90E+05 | 1.53E-02 | **5.23E-08** | ND |
| B8V32-siHPRT(A680) | hTfR1 | ND | ND | **6.65E-08** | ND |
| B8V32 | cTfR1 | 8.86E+05 | 3.86E-02 | **4.76E-08** | **4.21E-07** |
| B8V32-hFc-siSOD1 | cTfR1 | 1.04E+05 | 2.88E-02 | **2.83E-07** | **3.13E-07** |
| B8V32-hFc-siHPRT | cTfR1 | 1.04E+05 | 2.82E-02 | **2.75E-07** | **1.83E-07** |

ND: Not Determined. N/A: Not Applicable.
mTfR1, hTfR1, cTfR1: mouse, human or cynomolgus TfR1.

**Figure 2.**
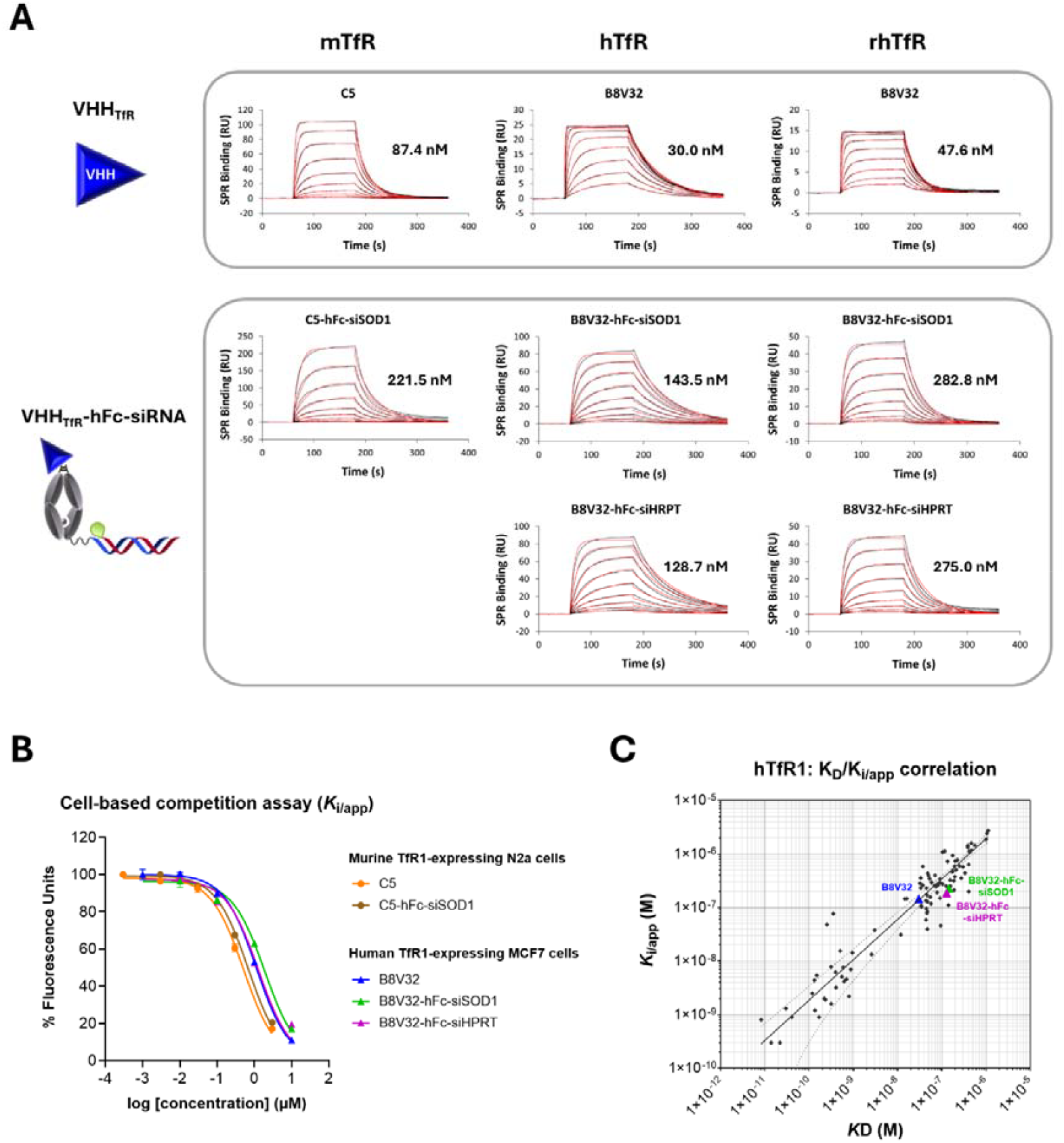
Cross-species TfR1-binding potential of free VHH and VHH-hFc-siRNA conjugates used in the present work. (A) Representative surface plasmon resonance (SPR) sensorgrams showing the association and dissociation profiles at pH 7.4 at increasing concentrations of unconjugated VHH or VHH-hFc-siRNA conjugates on immobilized murine, human or rhesus TfR (rhesus and cynomolgus monkey TfR share 100 % identity in their extracellular domain). RU, response units. Red traces correspond to fitting of experimental sensorgrams (black traces) using a Langmuir 1:1 binding model. (B) Apparent Ki on living murine TfR1-expressing Neuro-2a or human TfR1-expressing MCF-7 cells. Concentration-dependent competition curves obtained following 3 hrs co-incubation of indicated VHH or VHH-hFc-siRNA conjugates with the reference C5-A680 compound during 3 h at 37 °C. (C) Correlation between hTfR-binding potential as assessed using either SPR on immobilized hTfR-ECD (K_D_) or in the live-cell competition assay performed on human MCF-7 cells (Ki/app). Numerous compounds corresponding to VHHs from the same C5/B8 family (44), either unconjugated or conjugated to various payloads not described in the present study, were evaluated, allowing establishment of a regression line using GraphPad Prism v10 (solid line, Rsquare = 0.74, 95% confidence interval shown as dashed lines). Highlighted are the free B8V32 (blue) and B8V32-hFc-siRNA conjugates used in the present work (with siSOD1 in green and siHPRT in purple).

In addition to evaluating TfR1 binding in an immobilized format by SPR, we employed a complementary cell-based competition assay to determine the apparent affinity for a given epitope (K_i/app_) in living cells. While this assay integrates both cell-surface receptor binding and downstream endocytic processes, the strong correlation observed between SPR-derived *K*_*D*_ values and K_i/app_ values for hTfR1 supports its use as an integrated approach to assess TfR1 binding of VHH variants and conjugates in a physiologically relevant cellular context (**Figure 2C**). Accordingly, the C5-hFc-siSOD1 and B8V32-hFc-siRNA conjugates showed comparable K_i/app_ values for mTfR and hTfR, respectively, in the of range 200-400 nM (**Figure 2B**), confirming similar binding potential across species.

### Efficient TfR1-mediated delivery and release of siRNA in late endosomal compartments supports efficient RNAi activity in vitro

We first evaluated the intracellular trafficking profile of our VHH-siRNA conjugates and established a comprehensive picture of their functional delivery potential when conjugated to siRNAs. Alexa680, a highly stable and pH-insensitive dye with slow cellular elimination rate (□50h half-life) (57,58), was conjugated on the 3’-end of the antisense strand (AS) to allow direct and unbiased detection in subcellular compartments with no permeabilization and staining steps. These evaluations were performed in the human LN229 glioblastoma cell line using VHH-siRNA or VHH-hFc-siRNA conjugates with either subnanomolar (C5 or B8h1 conjugates) or moderate (B8V32 conjugates) affinity for hTfR1 (**Table 1**). Initial validation of their efficient and TfR1-dependent uptake in this cell line was obtained using the B8h1-siHPRT and B8V32-siHPRT conjugates, both showing strong and saturable uptake as compared to the C5neg-siHPRT control (**Figure 3A**). Consistent with their respective binding affinity to hTfR1, these conjugates showed uptake constants *K*_*m*_ of 6.1 and 104.6 nM, respectively. Based on these profiles, further intracellular and distribution studies were performed in the same cell line at the sub-saturating concentrations of 30 nM and 300 nM for high- and moderate-affinity conjugates, respectively. First, cellular elimination kinetics of different conjugates was investigated and compared to that of transferrin, using a previously described pulse-chase procedure (1-hour pulse followed by chase during indicated times in ligand-free medium) (58). All elimination profiles were robustly described by a two-phase decay regression, likely corresponding to an initial fast recycling phase along with TfR1, followed by slow elimination of non-recycled material from late vesicular compartments (**Figure 3B**). Contrary to transferrin that showed a typical fast and extensive initial decay (59-62), all TfR1-binding conjugates displayed very limited elimination after 6h chase, regardless of the binding affinity (high/sub-nanomolar affinity for C5- and B8h1-derived conjugates vs. moderate affinity for the B8V32-derived conjugate), the conjugate design (with or without Fc fragment), or the siRNA target. Second, investigation of the intracellular distribution of the moderate affinity B8V32-siHPRT conjugate showed that most of the siRNA payload was readily delivered to late endosomal compartments within a few hours, with □50% colocalizing with LysoTracker or LDL (low density lipoprotein) within a 3-hour incubation period (**Figure 3C-D**). In additional experiments, we verified that incubation of both high- or moderate-affinity conjugates did not alter TfR1-mediated uptake of transferrin (**Supplementary Figure 1C**). Taken together with the SPR data obtained at pH 6.0, this intracellular trafficking and distribution profile supports the hypothesis that the C5- and B8-derived VHH-siRNA conjugates are efficiently released from TfR1 within sorting endosomes, leading to substantial accumulation in late endolysosomal compartments.

**Figure 3.**
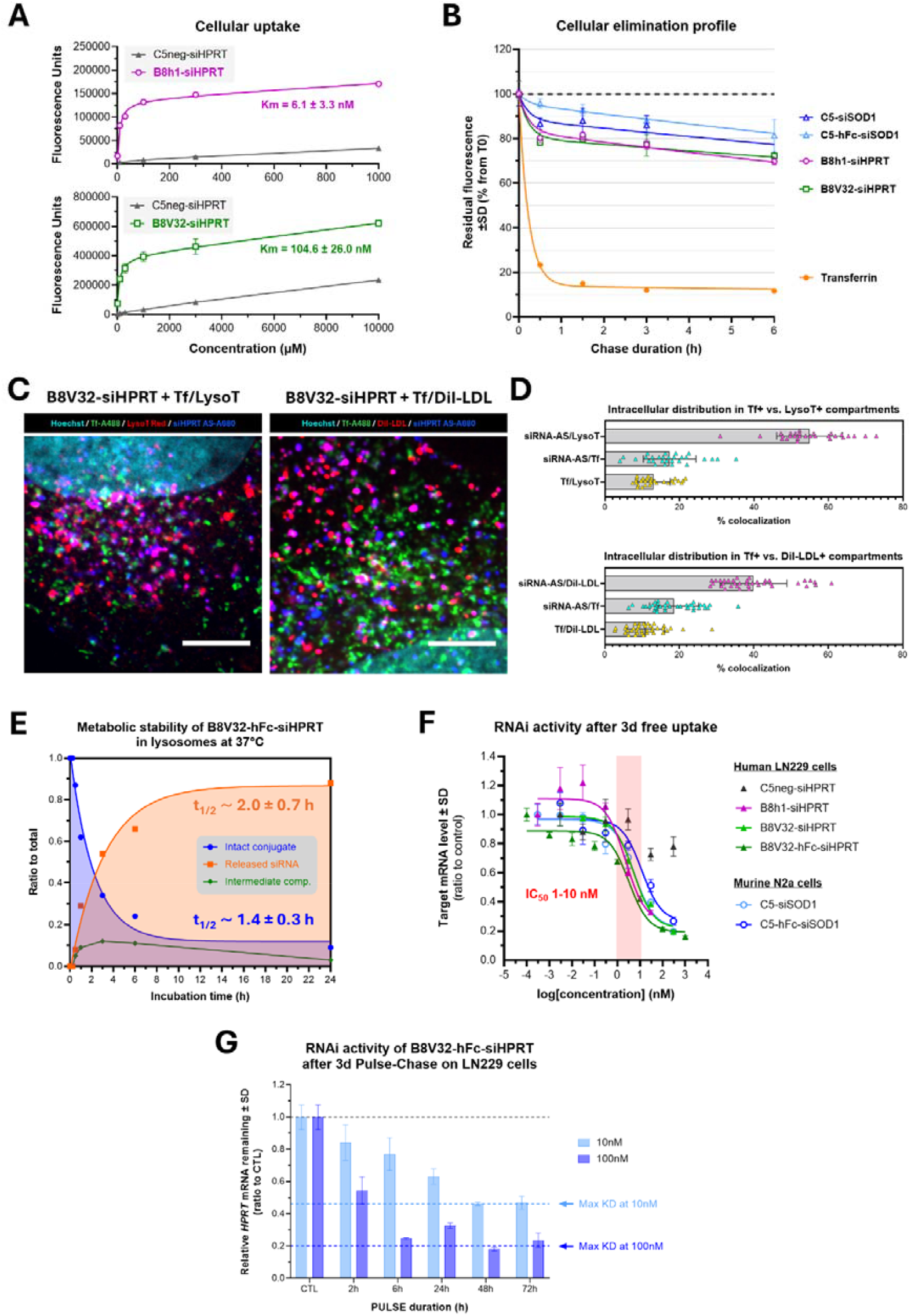
Intracellular delivery and target engagement of VHH-(hFc)-siRNA conjugates by free uptake in human and murine cells. (A) Human LN229 cells were incubated with increasing concentration of 3’AS-Alexa680 coupled VHH-siHPRT conjugates (with VHH = C5neg in dark grey, B8h1 in purple or B8V32 in green) during 3 hrs at 37 °C. The total A680-associated signal was then quantified on fixed cells using flow cytometry and shown as means ± SD. Km values were estimated by fitting experimental data using nonlinear regression. (B) Cells were pulsed during 1 h at 37 °C with the indicated A680-coupled VHH-(hFc)-siRNA conjugates (with siSOD1 or siHPRT) at sub-saturating concentrations (selected based on previous Km experiments: 300 nM for B8V32-siHPRT, 30 nM for C5- and B8h1-derived high affinity conjugates, 300 nM for C5neg-siHPRT), or Tf-A647 at 3 µg/mL, washed and then chased for the indicated times in ligand-free medium. The time-course of the total cellular-associated A680 signal on fixed cells was then quantified using flow cytometry. Results are expressed as means ± SD, normalized to the total signal measured at the end of the pulse (T0, set at 100%). (C) Shown are representative confocal micrographs of LN229 cells incubated during 3 hrs at 37 °C with the indicated A680-coupled VHH-siHPRT conjugates at sub-saturating concentrations. Late acidic compartments (i.e. late endosomes and lysosomes), early and recycling compartments were stained during the last hour by adding LysoTracker® Red DND-99 or DiI-LDL and Tf-A488, respectively, in the medium. Strong colocalization of A680-coupled siHPRT AS (false blue colour) with late compartments (false red colour) is observed in the perinuclear region (pink vesicular staining), with only minor colocalization with early and recycling compartments. Scale bars, 10 μm. (D) Quantification of intracellular distribution of A680-coupled siHPRT AS in either Tf-positive early and recycling compartments or in LysoTracker- or LDL-positive late endosomes and lysosomes. (E) Time-course of the metabolic degradation of the B8V32-hFc-siHPRT conjugate in rat liver tritosomes. Experimental data was generated from densitometry analysis of SYBR Gold pre-loaded 4% agarose gels (Supplemental Figure S1A); experimental curves showing the disappearance of the intact conjugate and appearance of the released siRNA were fitted using a nonlinear regression. (F) Target *Sod1* or *Hprt* mRNA knockdown after 72 hrs free uptake of indicated VHH-(hFc)-siRNA conjugates on murine Neuro-2a or human LN229 cells. Results are expressed as means ± SD and experimental data were fitted using nonlinear regression. Shown are representative experiments from at least n=3 independent experiments. (G) Human LN229 cells were incubated during indicated times with the B8V32-hFc-siHPRT conjugate at 10 or 100 nM (Pulse), followed by chase in ligand-free medium up to 72 hrs. Shown are target *Hprt* knockdown at the end of the 72 hrs Pulse-Chase period. Results are expressed as means ± SD.

We next examined the molecular fate of the conjugates following deposition in the acidic and proteolytically active environment of late endosomes and lysosomes. To this end, we used the B8V32-hFc-siHPRT conjugate, which was also selected for subsequent *in vivo* studies in hTfR1-expressing mice and NHPs. Upon incubation in lysosomal extracts, this conjugate exhibited rapid and extensive release of the siRNA payload, with a mean half-life of *ca*. 2.0 ± 0.7 h (**Figure 3E**). A slightly faster disappearance of the intact conjugate was observed, with a mean half-life of *ca*. 1.4 ± 0.3 h. This is consistent with the transient formation of an intermediate product which peaked at 3 hrs (based on its amount relative to total products in each lane) and still retained the siRNA moiety (**Figure 3E** and **Supplementary Figure 1A**). Interestingly, the Fc and VHH components of the conjugate exhibited comparable half-lives, with the Fc showing a slightly shorter and the VHH a slightly longer half-life (**Supplementary Figure 1A**). Moreover, introducing a PS linkage at the 3⍰ terminus of the siRNA resulted in a cleavage half-life similar to that observed with a PO linkage (**Supplementary Figure 1B**), indicating that the primary siRNA release mechanism is likely nuclease-independent. Instead, this initial cleavage most likely occurs at a site located between the amide bond formed during coupling of the heterobifunctional NHS-DBCO linker, and the Q-tag used for site-specific conjugation to the heterodimeric VHH-hFc precursor (**Figure 1B**).

Finally, different siHPRT conjugates were evaluated for their target mRNA knockdown (KD) potential by free uptake. These experiments were conducted in the same LN229 cell line, where the intracellular trafficking pattern was primarily investigated. Consistent with their comparable elimination kinetic profiles, all conjugates displayed similar TfR1-dependent RNAi activity, irrespective of the conjugation design and binding affinity to hTfR1, with IC_50_ values in the low nanomolar range (**Figure 3F**). Of note, conjugates used in previous intracellular trafficking experiments, comprising an Alexa680 in 3’AS, showed very similar RNAi activity (**Supplementary Figure 1D**). Similar results were also obtained using moderate affinity C5-derived conjugates on mTfR1-expressing murine Neuro-2a cells. These findings were best exemplified with the C5-siSOD1 vs. C5-hFc-siSOD1 conjugates, with mean IC_50_ values of 8.5 ± 7.8 nM and 6.3 ± 3.2 nM, respectively, and maximum KD values of 79.2 ± 6.8 % and 75.3 ± 5.8 %, respectively (**Figure 3F**). To investigate the actual exposure time needed to mediate meaningful endosomal depot, endosomal escape and target engagement, the B8V32-hFc-siHPRT conjugate was further evaluated for its RNAi activity using a pulse-chase approach. Consistent with the rapid intracellular delivery profile of these conjugates, we found that a 6-to 24-hour exposure time was sufficient to mediate robust KD after a 72-hour chase period (**Figure 3G**). This observation supports the hypothesis that short plasma exposure and efficient BBB-crossing may be sufficient *in vivo* to trigger efficient functional delivery in CNS parenchymal cells.

Collectively, C5/B8-derived VHH-siRNA conjugates showed efficient TfR1-mediated cellular uptake and intracellular delivery, translating into strong target engagement at low concentrations and with minimal exposure time. Considering the previously demonstrated CNS delivery potential of C5/B8-derived conjugates in mice (44), these features prompted us to further investigate their potential to trigger functional delivery of siRNAs in CNS tissues following systemic intravenous (IV) or subcutaneous (SC) administration in mice.

### IV or SC administration of a VHH-hFc-siSOD1 conjugate leads to comparable and robust functional delivery across brain regions in wild-type (WT) mice

CNS delivery of siRNA conjugates following systemic administration was first explored in WT mice using the moderate affinity conjugate C5-hFc-siSOD1 (K_D_ for mTfR1 of 222 nM). A single IV or SC administration at 1.5⍰mg/kg (siRNA molar equivalent) elicited a comparable 40% KD across all regions of the brain and spinal cord 14⍰days after dosing (**Figure 4A**). Further evaluation following single SC administration revealed very similar dose-response relationship across CNS regions, including deep regions such as the striatum, with maximal KD of the target mRNA ranging from 55 to 70% and ED_50_ values (dose eliciting 50% of the maximum KD effect) ranging from 0.57 to 1.24 mg/kg (mean 0.85 mg/kg siRNA) (**Figure 4B**). These data demonstrate not only efficient functional delivery of a *Sod1*-targeting siRNA across the BBB following low systemic dosing, but also excellent bioavailability when administered via the SC route. Consistent with the well-described saturability of TfR1-mediated transcytosis at the BBB (63), single IV at a 3-fold higher dose (4.5 mg/kg siRNA) did not lead to higher pharmacodynamic response in the CNS (**Supplementary Figure 2A**). This prompted us to compare the functional CNS delivery of the C5-hFc-siSOD1 conjugate following single SC administration at 4.5 mg/kg siRNA, approximately corresponding to the ED_80_, to the same total dose administered IV as a split-dosing regimen. Based on preliminary plasma PK studies with this conjugate (data not shown), the IV dosing regimen consisted of three low doses of 1.5 mg/kg siRNA administered every other day (IV Q2D x3, total dose 4.5 mg/kg siRNA). As expected, a single SC dose of 4.5 mg/kg siRNA resulted in a total plasma exposure over a 2-week period comparable to that achieved with the IV split-dosing regimen of 1.5 mg/kg siRNA, indicating excellent bioavailability using the SC route yet with a substantially smoother pharmacokinetic profile (**Supplementary Figure 2B**). In contrast, plasma exposure for both regimens was markedly lower than that observed with the non-binding C5neg-hFc-siSOD1 conjugate, consistent with rapid and extensive initial TfR1-mediated distributional clearance. Accordingly, both dosing paradigms produced similar levels of target engagement across CNS regions, whereas the non-binding control conjugate showed no detectable effect despite its substantially higher plasma exposure (**Supplementary Figure 2C**).

**Figure 4.**
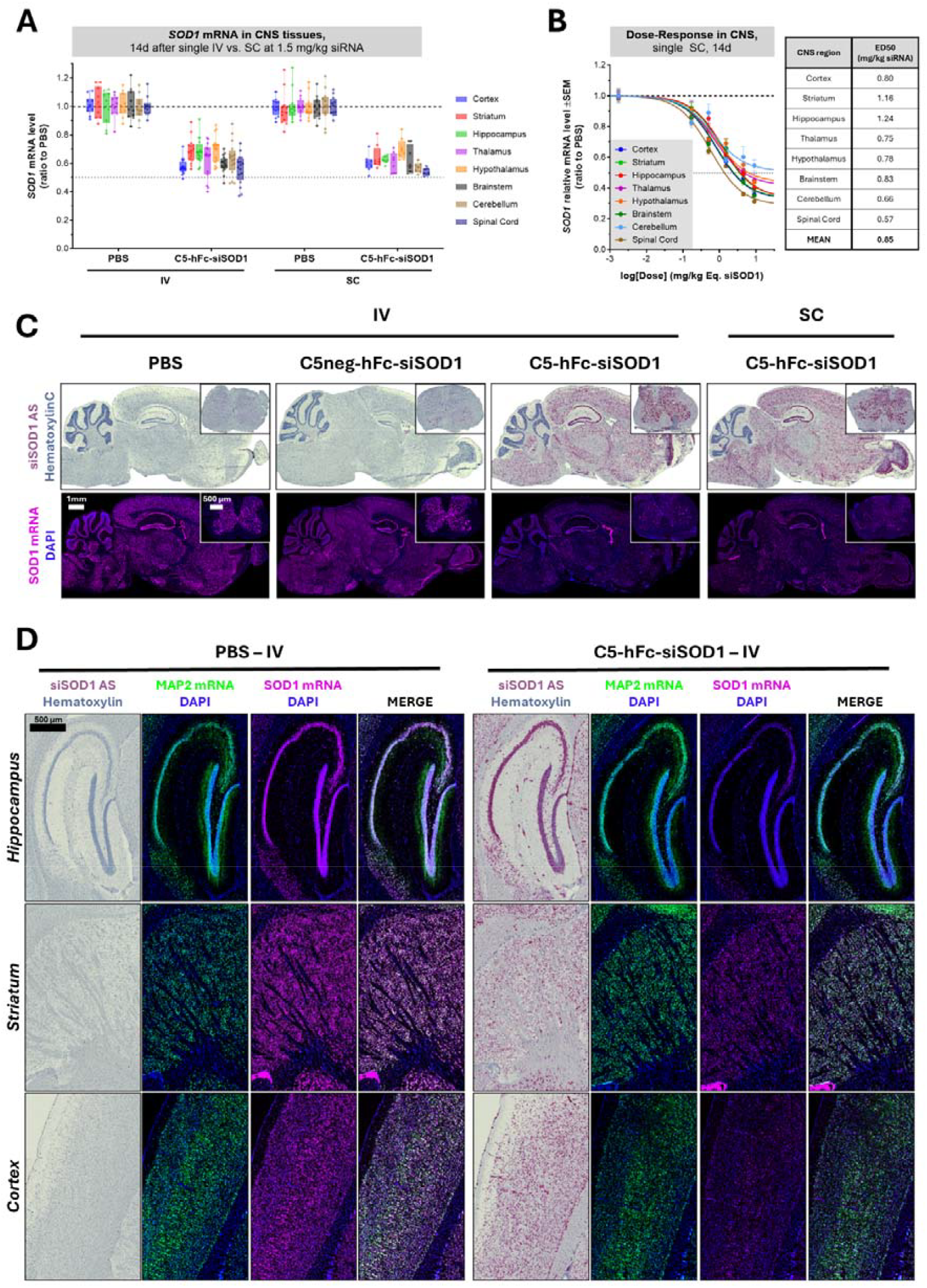
IV or SC administration of a mTfR1-targeting C5-hFc-siSOD1 conjugate allows broad distribution and target mRNA engagement throughout CNS regions in wild-type mice. (A, B) The C5-hFc-siSOD1 conjugate was dosed in WT mice as a single IV or SC bolus at 1.5 mg/kg (A) or 0.14 to 9 mg/kg siRNA equivalent dose (B). Target knockdown was measured 14 days after administration, using RT-qPCR on total RNA extracts from indicated regions of the CNS. The results are expressed as a ratio to mRNA levels measured in PBS-treated animals after normalization to RpL13 and RpL30 reference genes. (A) Box plots: extends from the 25th to 75th percentiles with center line plotted at the median. Whiskers indicate the smallest and the largest experimental values (n = 8-16 animals per group from 3 independent experiments). (B) Dose-response curves in each CNS region, where experimental results are expressed as means ± SEM, and estimated ED_50_ values. (C) Mice were administered with either multi-IV (Q2Dx3) at 1.5 mg/kg or single SC at 4.5 mg/kg siRNA equivalent dose of the C5-hFc-siSOD1 conjugate. Shown are *in situ* hybridization micrographs showing siSOD1 AS distribution (top panels, miRNAScope®) throughout the brain (sagittal sections) and spinal cord (cross-sections), and corresponding *Sod1* mRNA (lower panels, RNAScope®), 14 days after the last IV or single SC (representative animal from n=4 mice per group). (D) Magnifications and neuron-specific co-staining, using a Map2 mRNA-specific probe, in indicated regions of the brain.

These findings were further corroborated using *in situ* hybridization (ISH). Fourteen days after single SC or after the third IV dose, a widespread, TfR1-dependent depot of siSOD1 AS was detected predominantly within parenchymal cell bodies, with low signal in vascular-like structures, strongly supporting efficient BBB transcytosis and robust delivery to parenchymal cells. This distribution pattern correlated with pronounced KD of the target *Sod1* mRNA, which is primarily expressed in neurons positive for the pan-neuronal marker MAP2 (microtubule-associated protein 2), as illustrated in the hippocampus, striatum and cortex (**Figures 4C-D**). Notably, robust target engagement was confirmed across multiple, functionally distinct neuronal populations. These included substantia nigra dopaminergic neurons identified by the dopamine transporter (DAT) (**Supplementary Figure 3A**), striatal cholinergic interneurons identified by choline acetyltransferase (ChAT) (**Supplementary Figure 3B**), or ChAT-positive motor neurons in the facial nerve nucleus (cranial nerve VII, **Supplementary Figure 3C**) and in the ventral horn of the spinal cord grey matter (**Supplementary Figure 3D**). In contrast, no reduction in *Sod1* mRNA expression was detected in these neuronal populations following administration of the non-binding C5neg control conjugate (data not shown).

Collectively, these initial findings indicate that the favourable intracellular delivery profile and target engagement observed *in vitro* translates into rapid TfR1-mediated distribution in WT mice, resulting in widespread distribution and strong target engagement across all CNS regions following either IV or SC administration.

### Cross-species reactivity of our TfR1-binding VHHs enables efficient translation to hTfR1-expressing mice across multiple siRNA targets

Owing to the high degree of conservation of the TfR1 epitope recognized by the C5/B8 VHHs across rodents, NHPs, and humans, combined with the extensive and rational diversification of these parental hits (44), we were able to screen and identify suitable variants for subsequent assessment of siRNA functional delivery in hTfR1-expressing mice (data not shown). As shown in **Figure 2**, based on SPR, the B8V32-hFc-siSOD1 conjugate displayed matched TfR1-binding profiles compared to the C5-derived conjugate used in previous experiments in WT mice. Both conjugates are endowed with high on-rate together with fast off-rate and a resulting moderate binding affinity around 200 nM (**Table 1**). First, the plasma PK profile observed following multi-IV (Q2D x3) administration of B8V32-hFc-siSOD1 at 1.5 mg/kg siRNA was investigated and compared to the non-binding C5neg-derived conjugate, and to another B8V32-derived conjugate comprising a non-targeting control siRNA termed “siNTC” (**Figure 5A**, shown is the plasma PK profile after the last IV dose). As previously observed for the C5 variant in WT mice, both B8V32-derived conjugates exhibited similar rapid and extensive distributional clearance within the first 24 hrs. In contrast, the non-binding control conjugate showed only a modest (□1 log) decline, supporting a model of rapid and efficient TfR1-mediated tissue delivery. The plasma PK profile of the B8V32-hFc-siSOD1 conjugate was best described by a two-phase decay model, with an estimated clearance rate of 0.18 L/kg/d, an initial distribution half-life of 3.5 hrs associated with an approximately 2-log reduction within the first 24 hrs, and a terminal elimination half-life of ∼200 hrs. Importantly, quantification of the siSOD1 AS indicated stable association with the B8V32-hFc scaffold during the distribution phase. However, the terminal elimination phase was slightly more rapid for the siRNA than that of the molecular scaffold, likely reflecting progressive release of a plasma- or tissue-derived metabolite.

**Figure 5.**
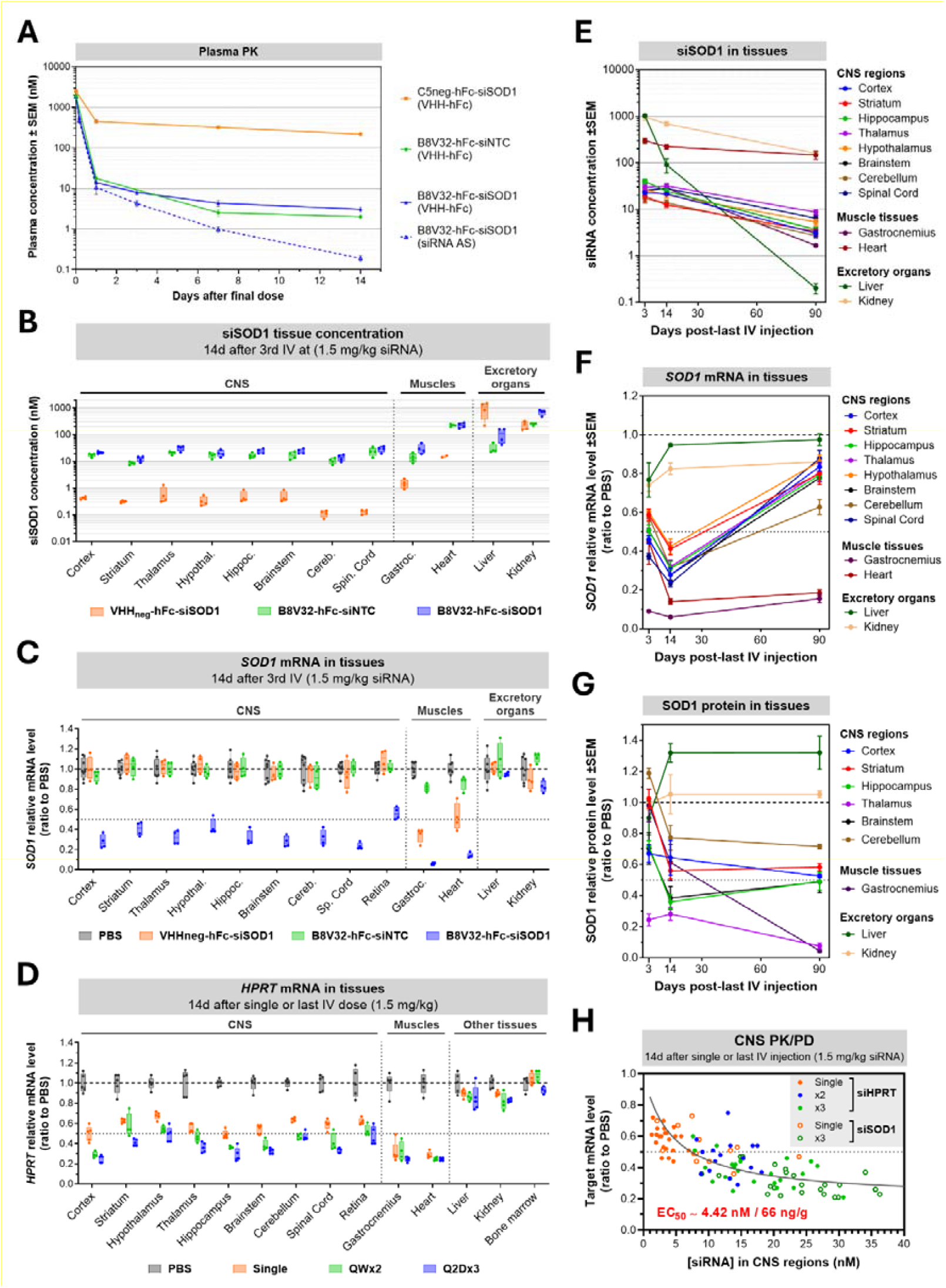
IV administration of hTfR-targeting B8V32-hFc-siRNA conjugates at low dose in hTfR1-KI mice achieve strong target engagement and PK/PD durability across brain regions. (A-C) Mice were dosed with the B8V32-hFc-siSOD1 conjugate at 1.5 mg/kg siRNA equivalent dose as multi-IV bolus (Q2D x3). (A) The plasma PK profile after the last IV administration was analyzed using either ELISA for measurement of the intact VHH-hFc scaffold, or stem-loop RT-qPCR for measurement of the intact siSOD1 AS. Shown are means ± SEM. Fourteen days after last IV dosing, tissues were processed to assess siRNA AS quantity (B) and target knockdown (C). The B8V32-hFc-siHPRT conjugate was administered as either a single or repeated IV bolus (QWx2 or Q2Dx3) at 1.5 mg/kg siRNA, and the target *Hprt* mRNA level were measured 14 days after single or last injection (D). (B-D) Box plots: extends from the 25th to 75th percentiles with center line plotted at the median. Whiskers indicate the smallest and the largest experimental values (n = 8 for PBS groups, n = 4 for treated groups). (E-G) Time-course of the tissue accumulation (E) and *Sod1* knockdown at the mRNA level (F) and at the protein level (G). Results are expressed as means ± SEM. (H) *Sod1* and *Hprt* mRNA silencing in CNS show a strong correlation with siRNA guide strand accumulation 14 days after single or repeated IV administration as indicated. Each plot represents individual value from one animal in one CNS region among the parieto-temporal cortex, striatum, hippocampus, thalamus, hypothalamus, brainstem, cerebellum or spinal cord. Experimental data were fitted with GraphPad Prism v10 using a three-parameter inhibitor vs. response regression model to estimate the EC_50_.

Consistent with a rapid TfR1-mediated BBB-crossing and tissue uptake of the B8V32-hFc-siSOD1 conjugate, a □100-fold higher accumulation of siSOD1 was found in CNS tissues 14 days after the last IV dose, as compared to the C5neg control conjugate (**Figure 5B**). Very similar concentrations were measured across all regions, ranging from 10 to 40 nM, suggesting efficient and homogenous BBB-crossing and deposition in parenchymal cells. At both early (3 days) and 14 days timepoints, CNS distribution of the siSOD1 AS closely mirrored that observed in WT mice with the C5 variant, with a widespread depot localized predominantly within parenchymal cell bodies and only minor localization in vascular-like structures (**Figure 6A**). These findings confirmed efficient engagement of the human TfR1 using the B8V32 variant. This early parenchymal depot was sufficient to drive robust and homogenous target engagement, with *Sod1* mRNA KD levels ranging from 60 to 80% across brain regions and the spinal cord at 14 days (**Figure 5C**). In contrast, although the B8V32-hFc-siNTC conjugate accumulated to similar levels in CNS tissues, it did not affect *Sod1* mRNA expression, confirming sequence-specific silencing by siSOD1. Importantly, a robust and specific □50% KD was also observed in the isolated retina. Further investigation using ISH confirmed TfR1-dependent uptake and depot in all layers of the retina (**Figure 6B**). The highest KD levels were observed in the retinal pigment epithelium (RPE), the photoreceptor inner segments, and the inner nuclear layer, which contains horizontal, bipolar, amacrine, and Müller cell bodies. Efficient functional delivery was also achieved in muscle tissues, with KD levels reaching up to 90% (**Figures 5B-C**). In these tissues, very low ED_50_ values were measured, ranging from 0.1 to 0.5 mg/kg siRNA in skeletal muscles and heart, respectively (data not shown). This finding contrasts with the very low TfR1 protein expression levels detected in skeletal muscle across species (**Supplementary Figure 4**). Nonetheless, it aligns with previous studies demonstrating efficient TfR1-mediated delivery of AOCs in these tissues (36,53,64), collectively suggesting that muscle tissues may be particularly prone to productive uptake of OTs. Accordingly, KD was also observed with the non-binding C5neg conjugate, though at levels significantly lower than those produced by B8V32⍰hFc⍰siSOD1.

**Figure 6.**
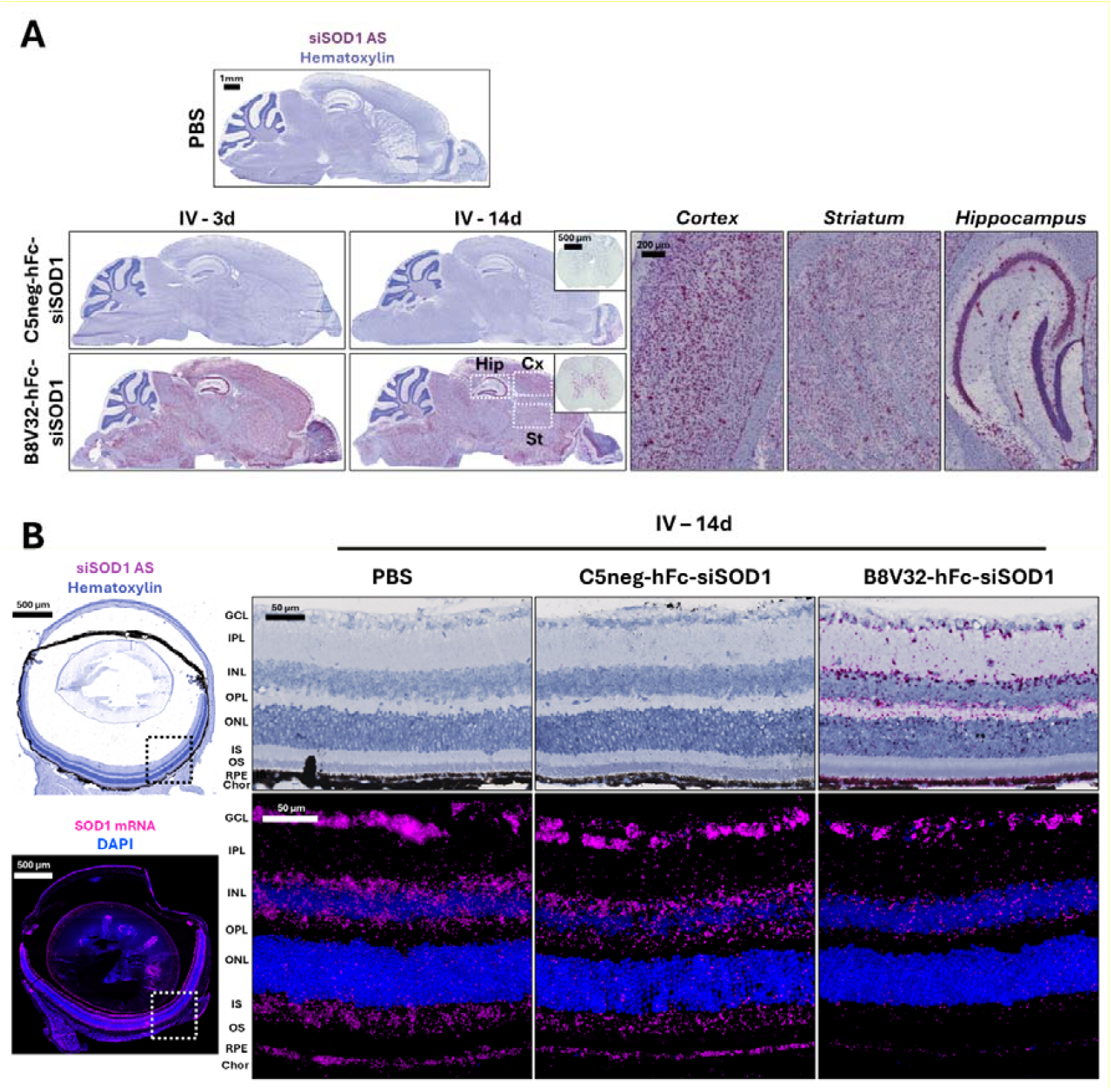
Brain and ocular distribution and target engagement of a B8V32-hFc-siSOD1 conjugate in hTfR1-KI mice following IV administration. (A-B) *In situ* hybridization micrographs showing siSOD1 AS distribution (miRNAScope®) throughout the brain (sagittal sections) and spinal cord (cross-sections) (A), 14 days after multi-IV bolus (Q2D x3) of the non-binding C5neg- or the TfR1-binding B8V32-hFc-siSOD1 conjugates, at 1.5 mg/kg siRNA. (B) *In situ* hybridization micrographs showing siSOD1 AS distribution (top panels, miRNAScope®) and corresponding *Sod1* mRNA (lower panels, RNAScope®) in retina layers of the same animals (representative animal from n=4 mice per group). Retina layers: GCL, Ganglion Cell Layer; IPL, Inner Plexiform Layer; INL, Inner Nuclear Layer; OPL, Outer Plexiform Layer; ONL, Outer Nuclear Layer; IS, Inner Segments; OS, Outer Segments; RPE, Retinal Pigment Epithelium; Chor, Choroid.

Another notable observation from the tissue PK/PD analysis was the lack of correlation between tissue accumulation and target engagement. This was exemplified by the heart and gastrocnemius muscle, with lower accumulation in the latter leading to a higher PD response, as well as by the liver and kidneys which exhibited the highest levels of tissue accumulation – likely reflecting their excretory roles – yet showed no measurable target engagement (**Figures 5B-C**).

A highly similar tissue distribution and PD profile was observed with another siRNA targeting *Hprt*, yielding 50-80% KD across CNS regions, approximately 80% KD in muscle tissues and minimal effects in excretory organs 14 days after the last dose (**Figure 5D** and **Supplementary Figure 5**). Although slightly higher siHPRT accumulation was detected in bone marrow compared with CNS regions or skeletal muscle (gastrocnemius), this increased exposure did not translate into detectable target engagement.

### CNS PK/PD relationship and duration of action in hTfR1-expressing mice

From a tissue PK standpoint, siSOD1 exhibited highly homogenous and stable accumulation across CNS regions. A mean siSOD1 concentration of 28 nM (corresponding to *ca*. 0.42 µg siRNA/g of tissue) was measured three days after the final IV dose (Q2D x3 at 1.5 mg/kg siRNA), with an apparent mono-exponential decay and an estimated half-life of nearly two months (54.2 days) (**Figure 5E**). Of note, these tissue concentrations are based on stem-loop qPCR-based quantification of the intact siRNA AS. Therefore, the total amount of potentially active metabolites resulting from 3’-trimming of the guide strand may be underestimated. A similar slow decay profile was observed for siSOD1 in muscle tissues and kidney. In contrast, liver exposure was only transient and decayed in parallel with plasma levels, indicating excretion rather than depot formation. In accordance with this rapid and sustained distribution profile of siSOD1 in CNS regions, a significant and homogenous target engagement was observed as of 3 days after the last dosing and persisted during at least 3 months (**Figures 5F-G** and **Supplementary Figure 6**). At this later time point, target KD remained substantial with 15-40% reduction at the mRNA level and 30-50% reduction at the protein level, consistent with the slower onset of protein suppression expected given the high stability of the SOD1 protein (65,66). Notably, this PD response was even more durable in muscle tissues, despite siRNA concentration half-lives comparable to those observed in the CNS. Given the similar *Sod1* mRNA expression levels in muscle and CNS tissues in basal condition (data not shown), this observation supports the hypothesis of a higher intrinsic KD efficiency in muscle. The CNS PK/PD relationship was further characterized using both siSOD1 and siHPRT conjugates after single, two (siHPRT only) or three IV doses. Although CNS exposure increased in a dose-proportional manner, as illustrated by the siHPRT accumulation profile (**Supplementary Figure 5**), the PD response reached saturation at tissue concentrations above ∼15 nM, with an estimated EC_50_ of 4.42 nM (corresponding to 66 ng siRNA/g of tissue) (**Figure 5H**).

Collectively, these findings, together with the *in situ* distribution data, demonstrate that the TfR1-targeting VHHs enable substantial and sustained accumulation of siRNAs within CNS parenchymal cells following low-dose systemic administration. This favourable CNS PK profile translates into a rapid onset and durable target KD, at both the mRNA and protein levels.

Overall, the robust CNS PK/PD relationship observed in mice strongly supports highly efficient TfR1-mediated transport across the BBB, and subsequent delivery of siRNAs to productive intracellular compartments within parenchymal cells.

### Systemic administration of VHH-siRNA conjugates enabled robust target engagement in muscle tissues and deep brain structures in NHPs

Initial translational validation from rodents to NHPs was obtained through an exploratory study conducted in olive baboons using a compact, □30 kDa C5-siSOD1 conjugate lacking a PK-enhancing moiety and incorporating a partially modified siSOD1 sequence (no 5’VP at the guide strand 5’-end). Because the *Sod1*-targeting siRNA used in mouse studies was rodent-selective, sequence mismatches were corrected to allow full complementarity of the guide strand with the ***SOD1*** mRNA from olive baboons (*Papio anubis*) (**Supplementary Figure 7A**). Despite substantial modifications in the seed region (4 nucleobase substitutions), the resulting “siSOD1h” retained silencing potency upon transfection in human cells when compared to the native siSOD1 on murine Neuro-2a cells, with an IC_50_ of □30 pM (**Supplementary Figure 7B**). Consistent with this high intrinsic potency, low nanomolar RNAi activity was also observed following free uptake of the C5-siSOD1h conjugate in multiple human cells lines, including MCF-7 and LN229 cells (**Supplementary Figure 7C**). Assessment of *in vivo* target engagement in olive baboons was restricted to muscle biopsies from the gastrocnemius, quadriceps and tibialis anterior. Following a single 30-min IV infusion at 5 mg/kg siRNA, robust and durable KD was achieved, with 50-60% *SOD1* mRNA KD across all muscle groups for up to 3-months, and with no clinical evidence of treatment-related adverse effects (**Supplementary Figure 7D**). Interestingly, a modest target KD was also detected with the non-conjugated siSOD1h. This effect may reflect the intrinsically high functional uptake capacity of muscle tissues, as suggested by our mouse data, and/or the presence of the 3’SS-aminohexyl modification which may confer increased lipophilicity. Collectively, these initial observations demonstrated that TfR1-mediated functional delivery of siRNAs can be achieved at low doses in a human-relevant species using VHH-based conjugates, enabled by a substantially smaller Ligand-siRNA architecture than that employed in previously described AOCs.

Because this preliminary study in olive baboons was not terminal, target engagement in CNS tissues could not be assessed. However, our initial mouse studies indicated that conjugates of this format require higher dose-levels to achieve meaningful functional uptake in the CNS (data not shown). Consequently, CNS delivery was subsequently evaluated in cynomolgus monkeys using the heterodimeric Fc-based design described in previous mouse studies. Given the inherent rodent/NHP cross-reactivity of the siHPRT tool and the ubiquitous expression of its target, together with the strong NHP/human cross-reactivity of the B8V32 VHH variant, we selected the same B8V32-hFc-siHPRT conjugate previously validated in hTfR1expressing mice. Cynomolgus monkeys received three weekly IV administrations (QW x3), at a dose of 1.5 mg/kg siRNA matching the dosing regimen used in the mouse studies. The plasma PK profile was best described by a two-phase exponential decay, characterized by a rapid and extensive initial distributional clearance (a 2-log decline within the first 2 days), similar to that observed in hTfR1 mice (**Figure 7A**). The estimated plasma clearance was 0.07 L/kg/d, which is slightly slower than that measured in mice (0.18 L/kg/d), with an initial distribution half-life of 6.4 hrs and a terminal elimination half-life of 49.7 hrs. No evidence of anti-drug antibody (ADA) formation was detected as comparable plasma concentrations were maintained following repeated administrations. Target engagement in tissues mirrored the profile observed in mice, with robust and homogenous KD across all brain regions. Specifically, *HPRT* mRNA reduction reached approximately 60% in cortical areas and up to 70% in deep brain regions, including the striatum (caudate and putamen) (**Figure 7B**). Importantly, no target engagement was detected in peripheral tissues, including excretory organs, except for muscle tissues where robust 80% KD was observed. CNS target engagement was further confirmed by ISH analysis, conducted on quarter ventral coronal sections of the left hemisphere at two anteroposterior levels: a rostral level encompassing the putamen (**Figure 7C**, to panels) and and a more caudal level encompassing the hippocampus (lower panels). In both regions, treated animals exhibited a pronounced reduction in *HPRT* mRNA expression across all cortical layers as well as comparably strong KD throughout deeper brain structures.

**Figure 7.**
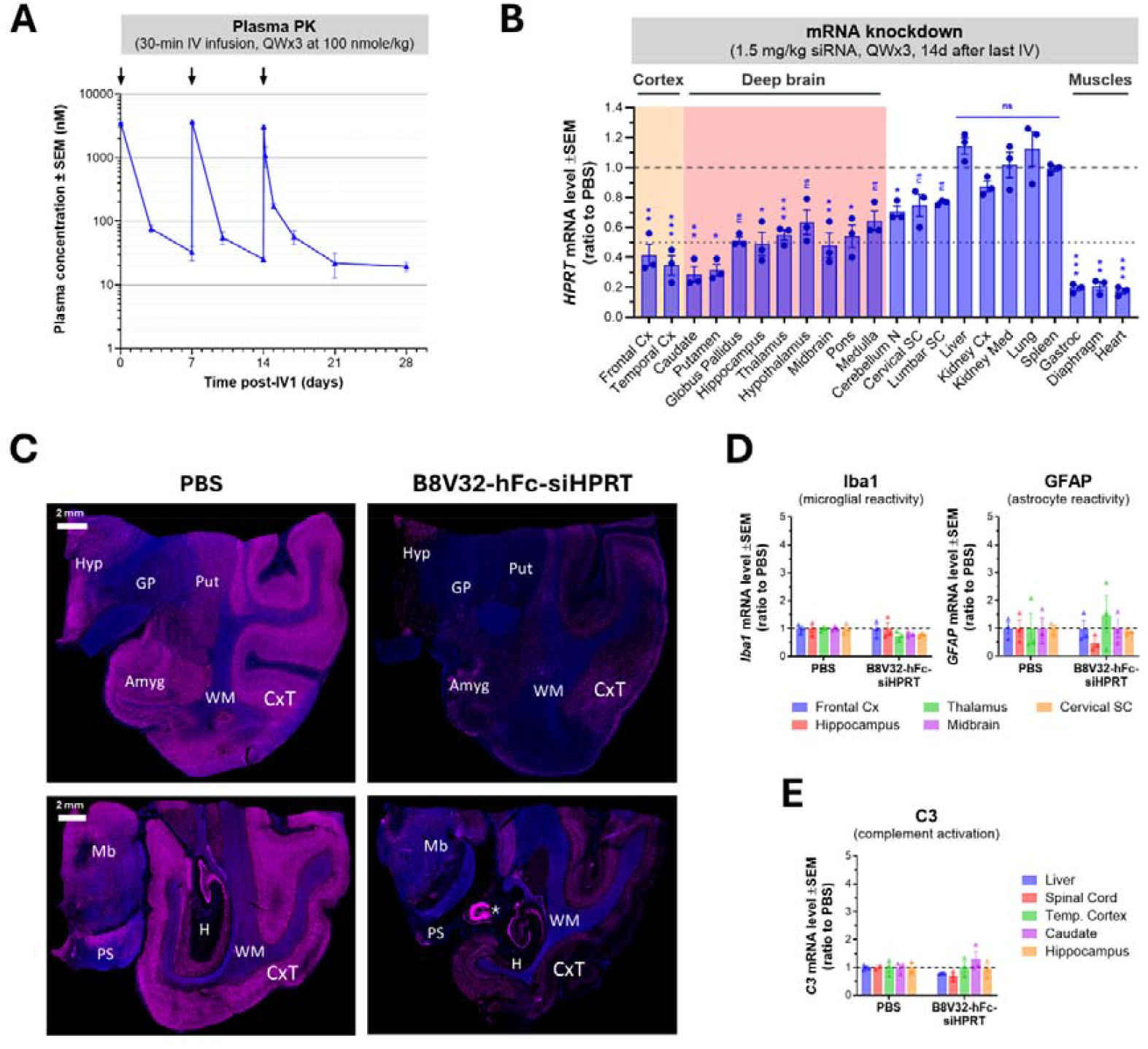
The B8V32-hFc-siRNA conjugate achieves strong target engagement in deep brain structures in non-human primates at low systemic dose. Cynomolgus monkeys were dosed by 30-min IV infusion three times at 7 days intervals with the B8V32-hFc-siHPRT conjugate, at the same dose previously evaluated in hTfR-KI mice of 1.5 mg/kg siRNA equivalent dose. Plasma samples were collected at indicated timepoints for PK analysis (A). Tissues were recovered at 14 days after last injection to assess target knockdown using RT-qPCR (B) and *in situ* hybridization in the brain using RNAScope® on one coronal section through the putamen (top panels: approximately -1 to -3 mm from the anterior commissure, based on Calabrese et al. 2015) (94) and one through the hippocampus (lower panels: approximately -11 to -13 mm from the anterior commissure; star indicates an artefact) (C). (D) Neuroinflammation was assessed on brain tissue samples by measurement of mRNA levels of IbA1, GFAP and C3 as markers of microglial reactivity, astrocyte reactivity and complement activation, respectively.

This tissue-selective target engagement profile was accompanied by a favourable safety profile. No adverse events were observed based on behavioural or clinical assessments, and neurological scores remained unchanged between treated and control animals. Haematology and blood chemistry parameters, including reticulocyte counts were unaltered (**Supplementary Figure 8**) and no systemic inflammation was detected based on longitudinal cytokine measurements. Liver and kidney functions were indistinguishable from those of control animals. Finally, assessment of neuroinflammation showed no evidence of microglial or astrocyte reactivity, nor any sign of complement activation across brain regions (**Figure 7D-E**).

## Discussion

Beyond the clinical success of ASGPR-targeted GalNAc–oligonucleotide conjugates for liver disease, several TfR1-targeting AOCs have now entered clinical development and show strong promise for the treatment of multiple muscle disorders (1,37,49). Owing to its high expression in the CNS, both at the brain endothelium and within parenchymal cells, TfR1 has long been regarded as an attractive target for deep-brain delivery of biologics. While this strategy has enabled the approval of a lysosomal enzyme replacement therapy and the advancement of multiple enzyme and antibody based programs, its translation to OTs lags considerably, despite substantial therapeutic potential (67). Several factors likely contribute to this gap. These include: i) the manufacturing complexity associated with conjugating biological targeting ligands to synthetic oligonucleotides (68); ii) the compounded safety liabilities arising from both components – such as perturbation of TfR1 physiological function alongside antibody- and oligonucleotide-mediated immunogenicity, inflammatory responses and off-target effects; and iii) the additional requirement for crossing the endosomal membrane to enable RNA target engagement. Bridging this gap therefore requires TfR1-targeting AOCs with optimized features across these dimensions, including scalable and robust manufacturing, low and transient systemic exposure, efficient BBB-crossing with productive uptake in parenchymal cells, and highly potent oligonucleotides with mitigated safety risks (69,70).

In the present work, we demonstrated that the TfR1-binding C5/B8 VHH platform displays attributes well suited to address these challenges, with strong translational consistency from rodents to NHPs. This platform drives broad CNS distribution of siRNAs leading to robust and durable target engagement following either IV or SC administration at clinically viable doses – namely 8.3 mg/kg of the full conjugate (1.5 mg/kg siRNA). For initial proof-of-concept studies, we leveraged the orthogonality, robustness and oligonucleotide compatibility of strain-promoted azide-alkyne cycloaddition (SPAAC), also referred to as click conjugation (52), to generate VHH-based AOCs with monovalent TfR1 binding and a defined Oligonucleotide-to-Antibody Ratio (OAR) of 1. Alongside alternative conjugation strategies – such as thiol-maleimide coupling to engineered cysteines (40,71,72) – this approach offers flexibility to tailor conjugate properties to specific target product profiles. Importantly, flexibility in conjugation chemistry is complemented by flexibility in conjugate design. In particular, we showed that compact VHH-siRNA conjugates of □30 kDa achieve robust and durable target engagement in muscle tissues following single systemic dose in both mice, with ED_50_ in skeletal muscles of 0.5 mg/kg siRNA (data not shown), and NHPs. These findings are in line with our *in vitro* observations of rapid and efficient trafficking of siRNAs to late endosomal compartments, independent of conjugate design. Although CNS delivery studies in this work primarily relied on Fc-based conjugates to extend plasma exposure, our data indicate that a transient systemic exposure window of 24 to 48 hours is sufficient to trigger robust and sustained CNS PD effects. This suggests that conjugate designs without the Fc may represent viable alternatives for certain applications.

One of the most distinctive features of the heterodimeric VHH-hFc-siRNA conjugates described here is their remarkably rapid and extensive distributional clearance across species, despite moderate TfR1 affinity. This pharmacokinetic profile, together with their adequate features enabling efficient BBB-crossing, correlates with robust target engagement in deep brain structures at low systemic dose. Based on our *in vitro* experiments, the mechanistic basis of this functional delivery profile likely resides in efficient TfR1-mediated productive uptake at low concentrations and with minimal exposure time. In contrast, conventional antibodies, other reported TfR1-targeting binding AOCs as well as our non-binding C5neg control exhibit markedly slower initial plasma decay, likely underlying slower or lower productive tissue uptake (40). Elucidating the molecular determinants underlying this distinctive PK behaviour is critical for rational optimization of CNS-directed AOC platforms. Our Structure-Activity Relationship (SAR) studies across C5/B8 variants revealed that even conjugates with sub-nanomolar TfR1 affinity — such as the C5-hFc-siSOD1 conjugate (0.68⍰nM; off-rate □10^−4^⍰s^−1^) — achieved efficient *in vitro* functional uptake and meaningful CNS target engagement in hTfR1-expressing mice (□30% KD, data not shown). Nevertheless, prevailing evidence suggests that moderate affinity and monovalent binding are optimal for TfR1-mediated crossing of the BBB (34,54,56,63,73-78). Recent studies further indicate that the specific TfR1 epitope targeted by a ligand critically influences intracellular trafficking of the ligand/TfR1 complex once internalized (33). Consistent with this concept, we observed that binding of our free or siRNA-conjugated VHHs to TfR1 decreased by 20-fold at the endosomal pH of 6.0, driven primarily by a marked 10-fold decrease in association rate, accompanied by lower binding responses. Structural modelling including docking experiments (data not shown) and prior work by David and colleagues indicated that our C5/B8 VHHs bind the TfR1 homodimer hinge region, contacting both receptor monomers (44). While histidine-mediated pH sensing is often implicated in endosomal dissociation (79,80), we did not identify histidine residues at the VHH-TfR1 interface. Rather, it has long been demonstrated that the TfR1 ectodomain, similar to the ASGPR or EGFR, undergoes major conformational changes when transitioning from plasma membranes at pH 7.4 to sorting endosomes at pH 6.0, leading to reduction of the exposed receptor surface (62). We therefore propose that such pH-dependent rearrangement of TfR1 ectodomain in sorting endosomes leads to mechanical dissociation of the C5/B8 VHHs. In turn, this would promote efficient release from TfR1 within brain endothelial cells and facilitate onward trafficking to late endosomal compartments in parenchymal cells.

Consistent with our *in vitro* experiments, our results in mice and NHPs indicate that transient systemic exposure at low dose is sufficient to establish high and sustained oligonucleotide depot throughout the CNS, resulting in a remarkably robust and durable PD response. Notably, target engagement in NHP was comparable or greater in deep brain structures relative to cortical regions. This distribution pattern highlights a key advantage of systemic OT delivery, which leverages the extensive and dense cerebral vasculature, over invasive local approaches such as ICV or IT administration. This PD profile may substantially enhance both efficacy and safety margins for OT candidates targeting diseases involving deep brain structures. Consistent with this premise, we observed robust target knockdown in dopaminergic neurons of the substantia nigra and in cholinergic interneurons of striatal regions, two structures centrally implicated in the pathophysiology of Parkinson’s disease (81,82). To assess functional delivery across CNS and peripheral tissues, we employed previously validated siRNAs targeting the ubiquitously expressed genes *Sod1* and *Hprt*. The use of siRNAs, rather than single-stranded ASOs, enabled an unbiased evaluation of TfR1-mediated uptake by avoiding confounding contributions from the intrinsic gymnotic activity of ASOs. However, within the brain, these mRNA targets are predominantly expressed in neuronal populations, which limited our ability to quantitatively assess functional delivery to other parenchymal cell types. Future studies using cell-type-specific siRNAs will be required to fully delineate the breadth of productive uptake across CNS cell populations. Beyond the brain, we also observed TfR1-mediated accumulation across all retinal layers, resulting in robust target engagement. This finding is consistent with the close structural and functional similarities between the BRB and the BBB, both of which protect specialized CNS compartments from circulating pathogens and toxins (19,20). These results suggest that our platform could provide a viable systemic alternative to invasive IVT administration for inherited retinal disorders, such as retinitis pigmentosa (RP) that affects photoreceptors, as well as acquired diseases including age-related macular degeneration (AMD) or diabetic retinopathy (83).

In hTfR1-expressing mouse models, VHH-hFc-siRNA conjugates displayed widespread, TfR1-dependent distribution across both CNS and peripheral tissues. Nevertheless, robust and durable target engagement was largely restricted to the CNS and muscle tissues. Notably, little or no KD was observed in the liver, kidney, or bone marrow, despite comparable or higher tissue siRNA exposure. This discrepancy between tissue distribution and productive uptake points to intriguing questions and potential hypotheses to explore. Interestingly, TfR1-targeting OTV conjugates incorporating a full-PS MALAT-1 gapmer ASO have been reported to induce relatively uniform 50% KD across CNS, muscle and several peripheral tissues in mice and NHPs (40). Whether this broader PD profile reflects a combination of TfR1-mediated delivery and non-specific, gymnosis-based uptake remains unresolved. Collectively, these observations suggest that individual TfR1-targeting AOC platforms may possess distinct strengths and limitations that must be aligned with the intended target product profile. Another notable finding from our studies is the lower ED_50_ and more durable PD response in muscle compared with CNS tissues, despite lower TfR1 protein expression in muscle and similar siRNA tissue exposure and decay kinetics. This striking PD profile in muscle does not appear to be specific to a particular oligonucleotide modality or delivery strategy, as comparable effects were reported for other TfR1-targeting and lipid-based platforms (53,64,84,85). Rather, we hypothesize that a modest, yet biologically meaningful increase in endosomal escape – one of the principal bottlenecks for productive OT delivery – may underlie this effect. This hypothesis could be tested by directly comparing target-engaged versus total intracellular oligonucleotide pools, using differential *in situ* imaging approaches.

From a safety perspective, the rapid distributional clearance and transient systemic exposure of our TfR1-targeting AOC platform may offer an additional advantage by limiting sustained engagement with immune and other liability pathways. Consistent with this notion both small VHH-siRNA conjugates and Fc-based formats were well tolerated in NHPs at the tested doses. No evidence of systemic inflammation, ADA formation, reticulocyte depletion, hepatic or renal dysfunction, or neuroinflammation was observed. Nevertheless, comprehensive safety assessments using therapeutic OT candidates and higher dose-levels will be essential to define safety margins and enable informed clinical translation.

Finally, the rodent/NHP/Human cross-reactivity of our C5/B8 VHHs — conferred by engagement of a highly conserved TfR1 epitope (44), provides two important advantages. From a regulatory standpoint, distinct C5 or B8 variants can be deployed as homologous or surrogate molecules across preclinical species, provided they maintain comparable binding properties. This principle is illustrated by the use of a C5-hFc-siSOD1 conjugate in WT mice, and a B8V32-hFc-siSOD1 conjugate in hTfR1-expressing mice and NHPs, both of which exhibited similar functional behaviour. Likewise, specific C5 variants may be deployed for efficacy and safety studies in NHPs, while alternative C5 or B8 variants can be selected for translation into humans. More broadly, diversification within the C5/B8 series enables adaptation to multiple oligonucleotide modalities, designs and chemistries, while preserving optimal TfR1-binding and BBB-crossing of the resulting AOC.

Taken together, our findings support a TfR1-targeting platform that combines robust functional delivery with substantial design flexibility, enabling the development of OT candidates across a broad spectrum of neurological and neuromuscular disorders. As BBB penetrant delivery technologies continue to mature — and as alternative BBB targets are emerging — it is increasingly clear that successful CNS drug development will require carefully matching the unique attributes of each delivery strategy to the specific biological and pathological context of individual diseases.

## Materials and methods

### Oligonucleotides synthesis and purification

siRNA sequences used in this work include siSOD1, targeting the murine superoxide dismutase 1 mRNA as previously described (5), a non-targeting control siNTC generated using AI-assisted design by Abzu (nucleobases permutations from the siSOD1 AS 2-21 sequence), a modified form of siSOD1, termed “siSOD1h”, targeting the human and olive baboons forms of the *SOD1* mRNA (**Supplementary Figure 7A**) and siHPRT, targeting the hypoxanthine-guanine phosphoribosyltransferase as previously described (36). All siRNAs consisted in 21/23 sense/antisense duplexes, with the following sequences and same stabilization schemes (“st23”): siSOD1 sense 5’-c•a•uuuuAaUCCucacucua•a•a-3’, antisense: 5’-VPu•U•uagAgUGaggaUuAaaaug•a•g-3’; siNTC sense 5’-u•a•accuUaAUCcucuacuu•a•a-3’, antisense: 5’-VPu•U•aagUaGAggauUaAgguua•a•g-3’; siHPRT sense 5’-u•c•cuauGaCUGuagauuuu•a•a-3’, antisense: 5’-VPu•U•aaaAuCUacagUcAuagga•a•u-3’, where lower-case letters indicate 2’-O-methyl (2’Ome) sugar modification, upper case letters indicates 2’-deoxy-2’-fluoro (2’F) sugar modification, • indicates phosphorothioate (PS) internucleosidic linkage and 5’VP indicates vinyl-phosphonate. For studies using siSOD1 in WT mice, nucleobases in positions 8 and 9 of the antisense strand were modified with a 2’F and no PS was added in 3’ of the sense strand (“st6”).

siRNAs were synthesized on 1-10 μmol scale on an K&A H-DNA/RNA/LNA Synthesizer using a 3’-Amino Modifier TFA Amino C-6 lcaa CPG 500Å solid support (54.8 μmol / g). The resin, the phosphoramidites and the reagents (diluent & washings) were purchased from Chemgenes (Wilmington, MA, USA). Fully protected nucleoside phosphoramidites were incorporated using standard solid-phase methods. ON synthesis conditions, i.e. 3 % trichloroacetic acid in dichloromethane (DCM) for deblocking, 0.25 M 5-(Ethylthio)-1*H*-tetrazole (ETT) in acetonitrile (MeCN) and 10% *N* -methylimidazole in tetrahydrofuran (THF) as activator; 10 % *N*-methylimidazole in THF, 10% acetic anhydride and 10 % pyridine in THF for capping; 0.05 M iodine in THF/pyridine/H2O (70:20:10) for phosphodiester oxidation; and 0.05 M 3-[(Dimethylaminomethylene)amino]-3H-1,2,4-dithiazole-5-thione (DDTT) solution in pyridine/acetonitrile for thiolation. All phosphoramidites were dissolved at 0.1 M in MeCN, addition of 10% DCM for 2’-O-methyluridine (mU), and incorporated with 12 min coupling times for all other phosphoramidites. After conclusion of the synthesis, the protecting groups were removed, and the ONs were cleaved from the resin by suspending the solid support in aqueous concentrated ammonia (30 %) and dimethylamine (3 %) at 65 °C for 5 hrs. The support was removed by filtration and the solution was concentrated using a speed-vac overnight at 65 °C. The crude ON was precipitated after addition of 200 mL H20 and 300mL AcONa (3 M) by using 1mL of cold ethanol. After centrifugation 10 min at 10000xg, the supernatant was removed, the oligonucleotide pellets were diluted in H20 and freeze-dried to obtain a white powder. The siRNA duplexes were obtained by hybridization of sense and antisense strand in Hepes buffer, at a final concentration of 200µM, after a heating period of 30 min at 95 °C and a cooling period with a first step at 37 °C and a second step at room temperature (RT). The hybridization was monitored by size exclusion chromatography. The identities and purities of all oligonucleotides were confirmed by electrospray ionization mass spectroscopy and reverse phase high-performance liquid chromatography.

### Generation, purification and characterization of VHH-(hFc)-siRNA conjugates

#### VHH-Myc-6His Production

Phagemids (pHEN1) encoding C5, B8V32, B8h1 and C5neg fused at their C-terminal to a Myc-tag and a hexa-histidine (6His) tag (final C-terminal sequence: AAAEQKLISEEDLNGAAHHHHHHGS) were transformed into *E. coli* BL21 (DE3) cells (ThermoFisher Scientific). Transformed bacteria were grown in 200 mL of 2YT medium until OD600 reached 0.5–0.8 and induced with 100 μM IPTG for overnight growth at 30 °C with shaking (250 rpm). Cells were pelleted and lysed by freeze-thawing combined with BugBuster® Protein Extraction Reagent (Novagen, Pretoria, South Africa). After centrifugation (4000xg, 20 min), VHHs were purified from the supernatant using metal affinity chromatography on TALON® Superflow™ resin (Cytiva, Marlborough, USA), following the manufacturer’s instructions. Bound proteins were eluted with 150 mM imidazole and desalted by diafiltration using 10 kDa MWCO Amicon® Ultra centrifugal concentrators (Merck, Darmstadt, Germany). An additional gel filtration step on a Superdex 75 column (Cytiva) combined with Mustang E (Pall, Port Washington, USA) filtration was performed to reduce endotoxin levels below 0.5 EU/mg. Endotoxin content was assessed using the Pierce Chromogenic Endotoxin Quant kit (ThermoFisher Scientific). Purified VHH-Myc-6His proteins were validated for purity and identity by SEC-UV and LC-UV-MS on an Orbitrap Exploris 240 (ThermoFisher Scientific).

#### Production of heterodimeric VHH-hFc-Myc precursors by transient transfection of Expi293TM cells

cDNAs encoding C5, B8V32, and C5neg VHH-hFc (derived from hIgG1) with LALA mutations (L234A/L235A, EU numbering), for studies in WT mice, or LALA-PG mutations (L234A/L235A/P329G, EU numbering), for studies in B-hTfR1 mice and NHPs, and knob mutation T366W (EU numbering), linked via a (G_4_S)_2_ linker between VHH and hFc, were obtained by gene synthesis, cloned into pcDNA3.4 (ThermoFisher Scientific) and fully sequenced. The cDNA encoding hFc-Myc with LALA-PG mutations and hole mutations T366S, L368A, Y407V (EU numbering) was also synthesized and cloned into pcDNA3.4 as hFcLALAPG-(Myc)1h. Heterodimeric VHH-hFc-Myc proteins were produced using the Expi293™ Expression System (ThermoFisher Scientific) by transient transfection with 1 µg total plasmid DNA per mL of culture, following the manufacturer’s protocol, using a 1:4 ratio of knob (VHH)1k-hFcLALAPG to hole hFcLALAPG-(Myc)1h plasmids. Cultures were harvested 72 h post-transfection, centrifuged (5 min at 300xg, then 15 min at 5000xg), filtered through 0.22 µm nylon membranes (Sigma-Aldrich), and purified on an ÄKTA Pure system using sequential steps: Protein A affinity chromatography (MabSelect™ PrismA resin, Cytiva), size-exclusion chromatography (SEC) (Superdex 200 column), and anion-exchange chromatography (AEX) (Resource Q 6 mL or HiTrap Q XL columns). Purified proteins were formulated in 20 mM Tris, 100 mM NaCl, pH 7.5, and validated for purity and identity by SEC-UV and LC-UV-MS. They were stored at -80°C until use for conjugation purposes.

#### Production of heterodimeric VHH-hFc-Myc precursors using a stable cell line

The generation of a stable cell line expressing B8V32-hFc-Myc was subcontracted to GTP Bioways (Labège, France). Briefly, two pCK expression vectors — (B8V32)1k-hFcLALAPG and hFcLALAPG-(Myc)1h — each carrying distinct antibiotic resistance markers, were co-transfected into the CHOvolution™ cell line (Celonic, Basel, Switzerland). Limiting dilution and screening identified a high-expressing clone, which was scaled up to 10 L in a bioreactor. Purification involved three steps: Protein A affinity capture, SEC and AEX. Final formulations were prepared in 20 mM Tris, 100 mM NaCl, pH 7.5, and validated for purity and identity by SEC-UV and LC-UV-MS and stored at -20°C until use for conjugation purposes.

#### Enzymatic Azide Functionalization of (VHH)1k-hFcLALAPG-(Myc)1h

Bacterial transglutaminase (BTG; Zedira, ref. T300) was resuspended in water according to the volume recommended in the manufacturer’s Certificate of Analysis. The amount of enzyme per vial (BTG units per vial) was used to calculate the stock concentration (units/µL). Aliquots were prepared to avoid repeated freeze–thaw cycles and stored at −20 °C or −80 °C. NH_2_-(CH_2_)_3_-N_3_ (Sigma-Aldrich) was used as azide donor. All reagents were of analytical grade. BTG-mediated functionalization was performed by adding NH_2_-(CH_2_)_3_-N_3_ (20 equiv.; 100 g/mol; density 1.02 g/mL) to (VHH)1k-hFcLALAPG-(Myc)1h (25 mg in 10.3 mL). BTG was then added at 0.0075 unit per nmol of protein (total 2.757 units; 19.9 µL). The final protein concentration in the reaction mixture was approximately 100 µM. Samples were mixed gently using pipette aspiration/dispensing; vortex mixing was avoided. Reactions were incubated at RT overnight. Reaction progress was monitored by LC–MS. Upon completion, samples were purified by size-exclusion chromatography (SEC) using an ÄKTA Pure system and a Superdex 200 26/600 GL column equilibrated in PBS. Fractions were analyzed by LC–MS; dilute fractions were concentrated using Amicon Ultra 15 centrifugal filters (10 kDa MWCO). Protein concentration was determined at 280 nm using a NanoDrop 2000 spectrophotometer (ThermoFisher Scientific). The resulting (VHH)1k-hFcLALAPG-(Myc)1h-N3 preparation was typically concentrated to 100-200 µM for subsequent conjugation.

#### Functionalization of siRNAs with DBCO-C6-NHS

Lyophilized siRNA (50 mg) was dissolved in 1.5 mL sterile water and stored at −20 °C as a stock solution. For functionalization, siRNA (1 equiv., 50 mg, 1.5 mL) was mixed with N,N-diisopropylethylamine (DIEA; 10 equiv.; 5.8 µL in water). The pH was adjusted to > 7. DBCO-C6-NHS ester (10 equiv.; 14.4 mg in DMF, total DMF volume 1.5 mL) was added to obtain a final H_2_0/DMF ratio of 1:1. The reaction was mixed briefly and monitored by LC–MS (RP-HPLC on a C18 column). Purification was performed using Amicon Ultra 15 centrifugal filters (10 kDa MWCO). Five wash steps were applied; the first used 20 % acetonitrile in water to prevent precipitation of unreacted DBCO-C6-NHS, followed by decreasing acetonitrile concentrations to finally exchange into PBS (pH 7.4). siRNA-DBCO concentration was quantified at 260 nm using a NanoDrop 2000 spectrophotometer (ThermoFisher Scientific). Preparations were stored at −20 °C.

#### Copper-Free Click Conjugation

DBCO–siRNA (1.3 equiv.; 5.7 mg in 438.2 µL PBS or water) was added to (VHH)1k-hFcLALAPG-(Myc)1h-N3 (1 equiv.; 19 mg in 2.41 mL PBS). The final protein concentration was maintained between 50 µM and 100 µM. Samples were gently homogenized by pipetting and incubated overnight at RT. Reaction monitoring was performed by SEC-HPLC. Final purification was carried out using SEC on an ÄKTA Pure system with a Superdex 200 26/600 GL column in PBS. Fractions were analyzed by SEC-HPLC; fractions with ≥ 95% purity were pooled. Diluted fractions were concentrated using Amicon Ultra 15 filters (10 kDa MWCO). Final conjugate concentration was quantified at 260 nm. The molar extinction coefficient of the conjugate corresponded to the sum of the coefficients of the protein and siRNA components; the 260 nm contribution of the protein was determined experimentally by measuring absorbance at 260 nm and 280 nm and applying the known 280 nm extinction coefficient from ProtParam. For optimal SEC resolution, injected volumes did not exceed 4% of the column volume (≤ 13 mL for a 320 mL column).

#### Characterization

All the final conjugates were characterized using 1) SEC-HPLC to assess their monomeric purity (Instrument: Thermofisher U3000 600bar; Column: AdvanceBio 300A 2.7um 7.8X300mm; Eluent : 0.15 M sodium phosphate; Flow rate: 1.2 mL/min; UV detection: 260 nm), 2) LCMS analysis to assess their identity (HPLC instrument : Thermofisher Vanquish; Column: Phenomenex Biozen 2.6 µm Oligo 100*2.1mm; Gradient: 2 % → 30% B from 1 to 13 min; Eluents : A = 12.5 mM HFIP, 4 mM DIEA in water; B = 12.5 mM HFIP, 4 mM DIEA in MeOH; Temperature : 75°C, Flow rate: 0.3 mL/min; UV detection: 260 nm UV-VIS 3; MS instrument: Thermofisher Exploris 240, negative electrospray mode) and 3) SDS-PAGE gel analysis used as a complementary analysis for purity assessment. Concentrations of the conjugates were calculated from absorbance at 260nm using a Nanodrop 2000 spectrophotometer (ThermoFisher Scientific).

### C5neg-hFcLALA-siSOD1mst6

Monomeric purity was > 99% - Deconvolution spectrum from the MS analysis allowed the attribution of the main peak to C5neg-hFcLALA-siSOD1mst6 sense strand, G0F_G1F glycosylation pattern (observed mass 75 133.95). The antisense strand is separated during the LC analysis from its sense strand. Concentration was calculated from the absorbance measured at 260 nm (ε= 419 100): 18.5 µM. SDS-PAGE analysis (one band) was in agreement with the expected molecular weight.

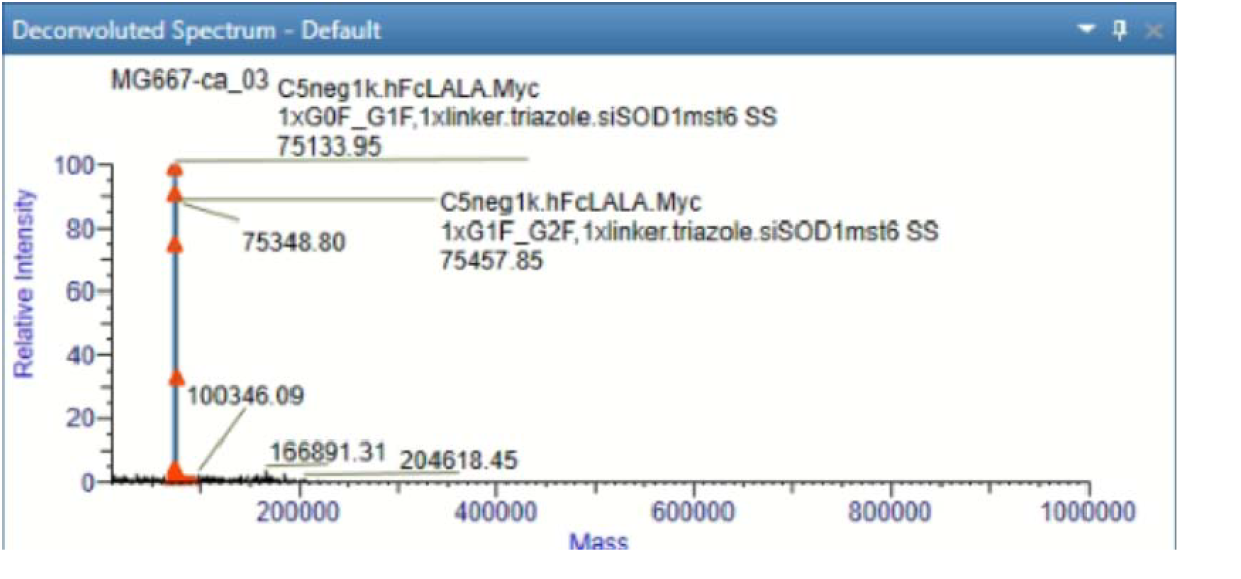

### C5-hFcLALA-siSOD1mst6

Monomeric purity was > 99% - Deconvolution spectrum from the MS analysis allowed the attribution of the main peak to C5-hFcLALA-siSOD1mst6 sense strand, G0F_G1F glycosylation pattern (observed mass 75 267.84). The antisense strand is separated during the LC analysis from its sense strand. Concentration was calculated from the absorbance measured at 260nm (ε= 419 100): 18.5µM. SDS-PAGE analysis (one band) was in agreement with the expected molecular weight.

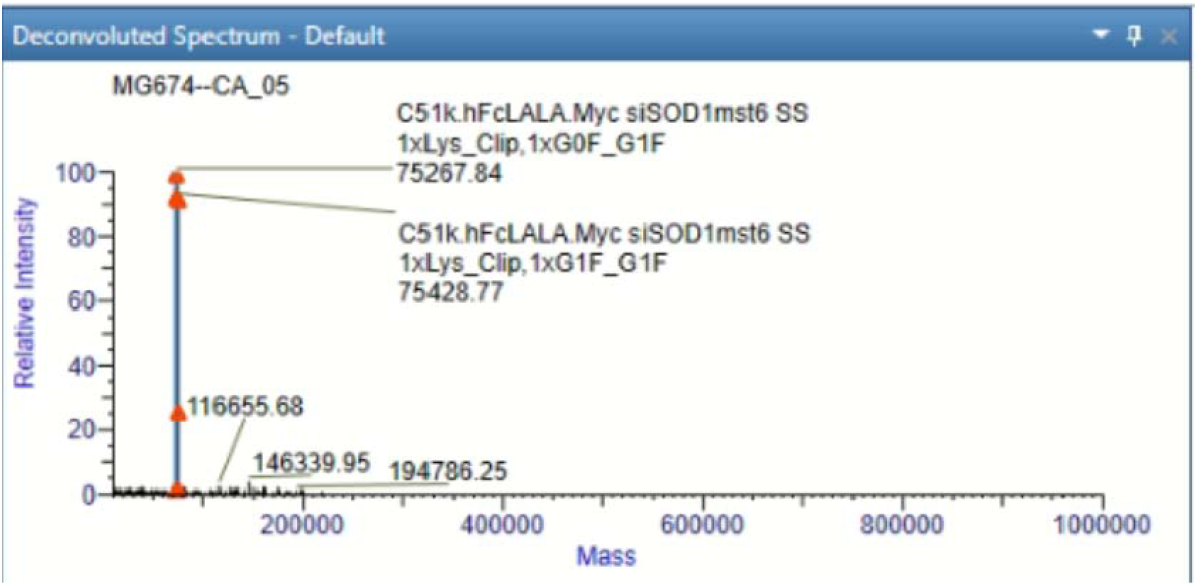

### C5neg-hFcLALA-siSOD1mst23

Monomeric purity was estimated to 98% - Deconvolution spectrum from the MS analysis allowed the attribution of the main peak to C5-hFcLALA-siSOD1mst23 sense strand, G1F_G1F glycosylation pattern (observed mass 75167.40). The antisense strand is separated during the LC analysis from its sense strand. Concentration was calculated from the absorbance measured at 260nm (ε= 467 860): 21.8 µM. SDS-PAGE analysis (one main band) was in agreement with the expected molecular weight.

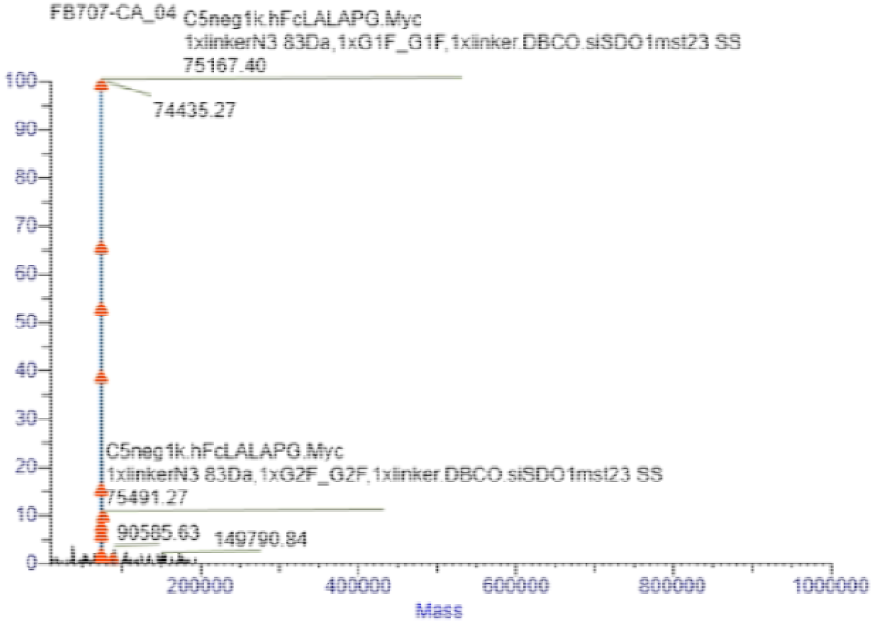

### B8V32-hFcLALA-siSOD1mst23

Monomeric purity was estimated to 97% - Deconvolution spectrum from the MS analysis allowed the attribution of the main peak to B8V32-hFcLALA-siSOD1mst23 sense strand, G0F_G0F glycosylation pattern (observed mass 75080.37). The antisense strand is separated during the LC analysis from its sense strand. Concentration was calculated from the absorbance measured at 260nm (ε= 471 925): 21.43 µM. SDS-PAGE analysis (one main band) was in agreement with the expected molecular weight.

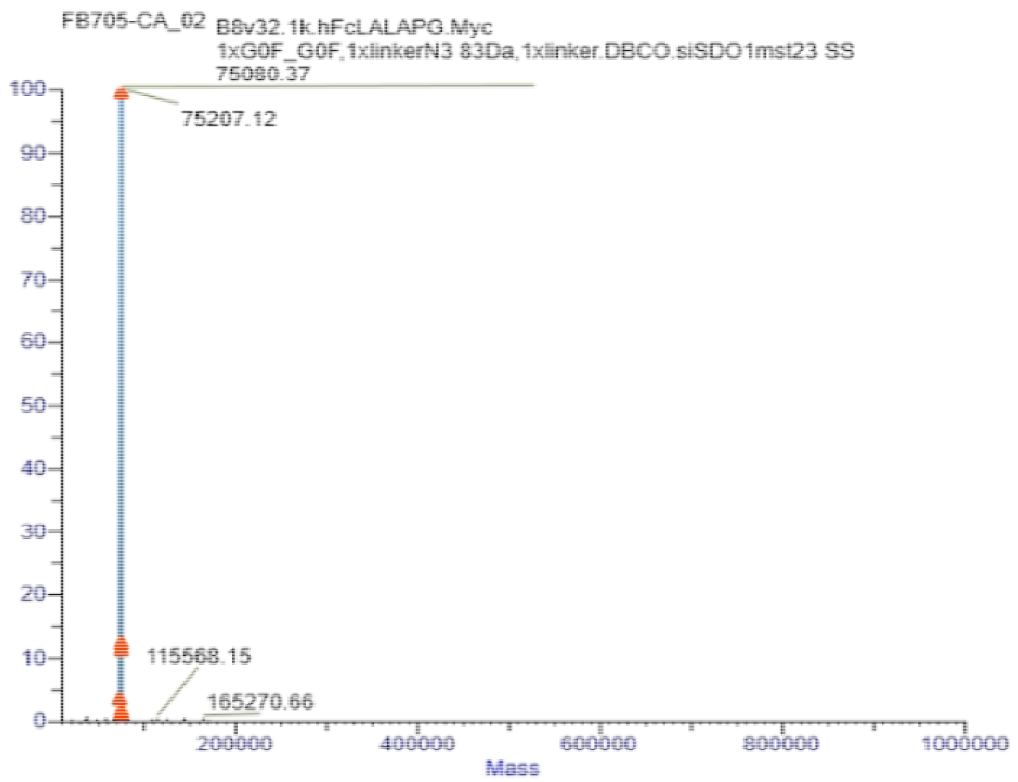

### B8V32-hFcLALAPG-siHPRT

Monomeric purity was estimated to 89% - Deconvolution spectrum from the MS analysis allowed the attribution of the main peak to B8V32-hFcLALA-siSOD1mst23 sense strand, G0F_G0F glycosylation pattern (observed mass 75176.67). The antisense strand is separated during the LC analysis from its sense strand. Concentration was calculated from the absorbance measured at 260nm (ε= 473 173): 20.9 µM. DS-PAGE analysis (one main band) was in agreement with the expected molecular weight.

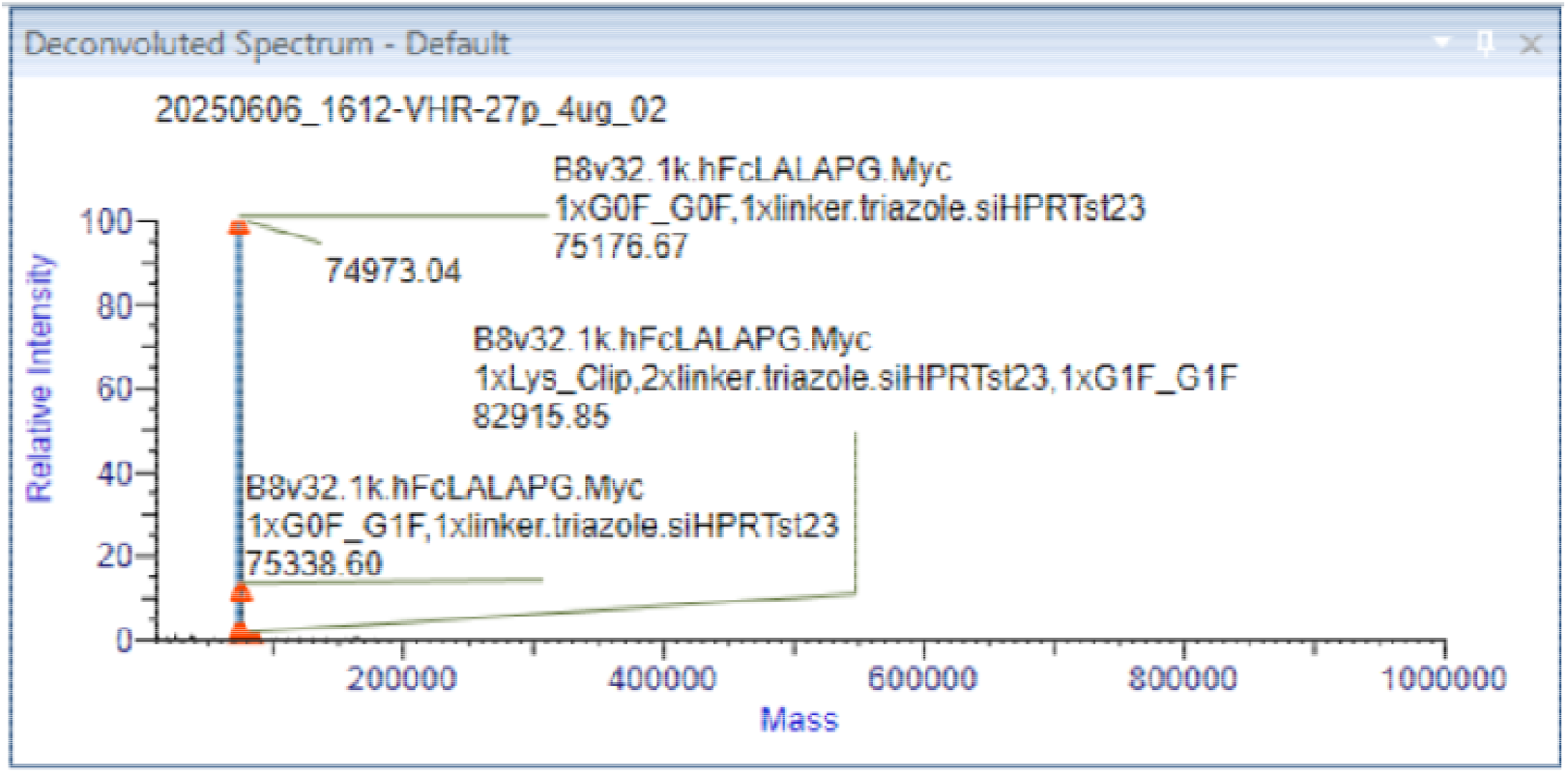

### Determination of TfR1-binding affinity of VHH-(hFc)-siRNA conjugates using Surface Plasmon Resonance

SPR measurements were performed at 25 °C using a Biacore T200 apparatus (Cytiva) and 50 mM HEPES-NaOH pH 7.4, 150 mM NaCl 0.005% Tween-20 (v/v) as running buffer. The N-terminal end of the ectodomain (ECD) of human TfR1 (hTfR1), mouse TfR1 (mTfR1) and rhesus/cynomolgus monkey TfR (noted “cTfR1”; GeneBank references NM_003234.2, NM_011638 and NC_041755.1, respectively) were fused to the C-terminal end of a mouse IgG2b Fc fragment and the recombinant proteins were produced and purified in-house. Of note the ECD of TfR1 from rhesus (***macaca mulata***) and cynomolgus (*macaca* fascicularis) monkeys are 100% identical. For SPR experiments, an anti-mouse antibody (Sigma Alrich) was first immobilized via amine coupling at a density of 100 fmol/mm2 onto a CM5 sensorchip (Cytiva), then the mFc-hTfR, mFc-rhTfR or mFc-mTfR ECD ligand was immobilized by affinity interaction at a density of 4.8 - 9.7 fmol/mm2. A control flowcell with anti-mFc antibody but without immobilized receptor was used as reference. VHH and VHH-(hFc)-siRNA conjugates were prepared by two-fold dilutions (range 5-1280 nM) in running buffer and injected over the flowcells during 120 s at 30 μL/min with a dissociation time of 200s. Blank runs of running buffer were performed in the same condition and subtracted from sample runs before evaluation. Double-subtracted sensorgrams were globally fitted with the Langmuir 1:1 binding model from Biacore T200 Evaluation version 3.2. Data are representative of at least 3 independent experiments.

### Competition assay in live cells

This cell-based competition assay was initially developed to evaluate the binding potential of peptides and peptide-cargo conjugates targeting the LDL receptor (86), by determining the apparent inhibition constant (Ki/app) values from experimental inhibition curves obtained in living cells. Here, it was adapted to evaluate the binding potential of unconjugated VHH or VHH-(hFc)-siRNA conjugates, derived from the C5/B8 VHH variants, on living TfR-expressing murine Neuro-2a (N2a) or human MCF-7 cells. Briefly, N2a and MCF-7 cells were grown in Dulbecco’s Modified Eagle Medium (DMEM) GlutaMAX™ supplemented with 10% v/v FBS and 100U/mL Penicillin and 100µg/mL Streptomycin in 5% CO2 at 37°C. Cells were seeded two days before experiments in 96-well plates at a density of 40.000 cells/well for N2a cells or 30.000 cells/well for MCF-7 cells. On the day of the experiment, the cells were co-incubated with increasing concentrations of free VHHs or VHH-hFc-Oligo conjugates and a sub-saturating concentration of the fluorescent reference compound C5-Alexa680 (100 nM for N2a cells or 10 nM for MCF-7 cells) diluted in DMEM supplemented with 1% bovine serum albumin (BSA) during 3 hrs at 37°C. At the end of the incubation period, the total cellular A680-associated fluorescent signals were acquired and analyzed as described in the ‘Flow Cytometry’ section. GraphPad Prism® software v10 was used to determine the apparent K_i_ inhibition constants using nonlinear regression.

### Evaluation of endocytic constant Km and determination of sub-saturating working concentrations

The endocytic constant *K*_*m*_, representing the concentration allowing binding and endocytosis of the tested compound at the half-maximal capacity of the cellular model, was evaluated in human glioblastoma LN229 cells. Cells were grown in Dulbecco’s Modified Eagle Medium (DMEM) GlutaMAX™ supplemented with 10% v/v FBS and 100U/mL Penicillin and 100µg/mL Streptomycin in 5% CO2 at 37°C. Two days before the experiment, cells were seeded in 96-well plates at a density of 30.000 cells/well. On the day of the experiment, the cells were incubated with increasing concentrations of siRNA 3’-AS A680-coupled VHH-(hFc)-siRNA conjugates diluted in DMEM supplemented with 1% bovine serum albumin (BSA) during 3 hrs at 37°C. At the end of the incubation period, the total cellular A680-associated fluorescent signals were acquired and analyzed as described in the ‘Flow Cytometry’ section. GraphPad Prism® software v10 was used to determined *K*_*m*_ values using nonlinear regression. Sub-saturation working concentrations for further intracellular trafficking experiments were set at *ca*. 3-fold the *K*_*m*_ value.

### Evaluation of cellular elimination

Total cellular elimination of A680-coupled VHH-(hFc)-siRNA conjugates was studied using a pulse-chase protocol (1-hour pulse at 37 °C followed by chase in ligand-free medium during 30 min, 1.5 hrs, 3 hrs, 3 hrs or 6 hrs at 37 °C). LN229 cells were grown as described hereabove and seeded in 96-well plates at a density of 30,000 cells/well two days before the experiment. Each A680-coupled VHH-(hFc)-siRNA conjugate was incubated at sub-saturating concentration, based on previous determination of individual *K*_*m*_ values. At the end of the pulse (T0) or after indicated chase times, the total cellular A680-associated fluorescent signals were acquired and analyzed as described in the ‘Flow Cytometry’ section. The residual amount of A680-coupled VHH-(hFc)-siRNA conjugates at the end of the pulse-chase procedure was calculated as a ratio to the total A680-associated signal measured at the end of the pulse (set at 100%). GraphPad Prism v10 was used to identify fitting curves that best described individual experimental datasets. For all elimination profiles, the plateau value was set to zero as the A680 fluorophore is assumed to undergo complete cellular elimination over time.

### Evaluation of steady-state intracellular distribution

LN229 cells were grown as described hereabove and seeded onto glass coverslips in 24-well plates at density of 10,000 cells/well two days before experiment. On the day of the experiment, the cells were incubated for 3 hrs with sub-saturating concentrations of fluorescent VHH-siRNA conjugates with DiI-LDL (L3482, ThermoFisher Scientific) at 100 µg/mL or with LysoTracker Red DND-99 (only during the last hour) at 1:10,000 diluted in DMEM supplemented with 1% bovine serum albumin (BSA) at 37°C in 5 % CO2 at 37 °C. At the end of incubation, cells were washed 3 times in cold D-PBS and then fixed by 4% PFA for 10 min at RT. Nuclei were stained with Hoechst 1:1000 solution. Coverslips containing cells were mounted on glass slides in Prolong Gold after washing 3 times in PBS. Images were acquired using a Nikon Ti2 inverted microscope equipped with a Yokogawa Nipkow CSU-W1 Nipkow confocal disc head and the SoRa resolution enhancement module with 60x objective and X4 lens. JACoP plugin of ImageJ software was employed to quantify the intracellular distribution of A680 in either Tf-A488-positive (early, sorting, and recycling endosomes) or DiI-LDL-positive / LysoTracker Red DND-99 positive (late endosomes and lysosomes) compartments. An average of 35 images were analyzed for each group.

### Transferrin uptake assay

LN229 cells were grown as described hereabove and seeded in 96-well plates at density of 15,000 cells/well two days before experiment. On the day of the experiment, cells were incubated with sub-saturating concentrations of fluorescent VHH-siRNA-A680 conjugates, diluted in DMEM supplemented with 1% bovine serum albumin (BSA) at 37°C for 3 hrs in 5 % CO2 at 37 °C. Then, cells were washed with DPBS and incubated with 3 µg/mL of Tf-A88 (T13342, ThermoFisher Scientific) during 1 h in 5 % CO2 at 37 °C. At the end of the incubation period, the total cellular A488 and A680-associated fluorescent signals were acquired and analyzed as described in the ‘Flow Cytometry’ section.

### Flow cytometry

At the end of live-cell procedures, cells were extensively washed with D-PBS and dissociated in Trypsin/EDTA 0.05% for 5 min at 37 °C before addition of cold complete culture media (4 °C) to inhibit Trypsin activity. Resuspended cells were transferred in a 96-deepwell plate (V-bottom) containing 1 % fetal bovine serum (FBS) / 0.02% sodium azide / 5 mM EDTA in phosphate-buffered saline (PBS) and then centrifuged for 3 min at 700xg at 4 °C. After removing the supernatant, living cells were stained using LIVE/DEAD Violet reagent (ThermoFisher Scientific) diluted at the appropriate concentration in cold PBS and incubated for 30 min at 4 °C in the dark. Cells were then washed by addition of 550 µL of cold PBS and then centrifuged for 3 min at 700xg at 4°C. After removing the supernatant, cells were fixed in D-PBS containing 5 mM EDTA and 4 % paraformaldehyde (PFA) (v/v 1:1) for 15 min at RT in the dark. PFA was diluted by addition of 500 µL of PBS, then removed by centrifugation (3 min, 700xg, 4 °C) and cells were resuspended in D-PBS containing 5 mM EDTA. Then the cellular A488- and/or A680-associated fluorescence signal was acquired using an Attune™ NxT flow cytometer (ThermoFisher Scientific) equipped with Attune™ NxT v3.1.2 following MiFlowCyt recommendations (87,88). Data were analyzed using the FlowJo software v10.10.0.

### Metabolic stability in rat liver tritosomes

Rat liver lysosomal extracts (tritosomes) were used as a predictive *in vitro* test system of lysosomal stability. This assay was used here to evaluate the lyososomal metabolic stability of VHH-(hFc)-siRNA conjugates used in the present work and was adapted from the method described in Weingärtner et al., 2020 (89). Briefly, Sprague-Dawley rat liver tritosomes (X000035, Tebubio) were thawed to RT then placed in an acidic buffer (1.5 M acetic acid, 1.5 M sodium acetate, pH 4.75) in 10:1 proportion to mimic the acidified environment. Thirty µL of acidified tritosomes were added to 10 µL of VHH-(hFc)-HPRT at 20 µM and incubated at 37 °C for 5 min, 15 min, 30 min, 1 hr, 3 hrs, 6 hrs, 24 hrs and 7 days. Samples were analyzed using agarose gel electrophoresis for RNA detection, Western-Blot for VHH-hFc detection (see Western-Blot section), and stem-loop RT-qPCR for siRNA AS quantification. For RNA detection, 150 ng siRNA equivalent of each time point were loaded into E-gel EX 4 % agarose gel (G401004, ThermoFisher Scientific). TrackIt™ Ultra Low Range (10488023, ThermoFisher Scientific) was used as a DNA Ladder. The gel was placed into E-Gel™ Power Snap Plus (ThermoFisher Scientific) and ran for 15 min. At the end of the migration, the gel was imaged using E-Gel™ Power Snap Camera (ThermoFisher Scientific).

Gel’s images were quantified using ImageJ’s software (v1.54g): first an average background subtraction was applied to the images excluding the lane’s labels. To improve image contrast and correct for potential baseline drifts on averaged lane’s signals, we Z-scored the background-subtracted images pixel by pixel as follows:

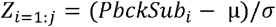

where Zi is the resulting Z score pixel value (i), j the total number of pixels, PbckSubi the background subtracted pixel value, µ and σ are respectively the mean and standard deviation of PbckSub for all pixels. The resulting images are thus directly expressed as a statistical measure (standard deviation compared to the mean). The outcome also facilitates manual detection of molecular weight peaks using the imageJ gel’s tools (plot lanes). The resulting area under the curves (AUCs) were then extracted for each peak contained in a gel’s lane to finally compute the PK profile using Graphpad Prism Software v10. One phase decay nonlinear regression was used to best describe individual experimental datasets and estimate half-lives.

### Target mRNA knockdown by free uptake

Murine neuroblastoma Neuro-2a (N2a) cells and Human glioblastoma LN229 were grown in DMEM GlutaMAX supplemented with 10 % v/v FBS and 100 U/mL Penicillin and 100 µg/mL Streptomycin in 5 % CO_2_ at 37 °C. For free uptake experiments, cells were seeded in 96-well plates at a density of 3,000 cells/well for N2a and 5,000 cells/well for LN229. Twenty-four hours later, the medium was carefully removed, and cells were further incubated during 3 days in DMEM supplemented with 1 % v/v FBS containing increasing concentrations of conjugated oligos. Cells were then washed with D-PBS and RNA extraction and reverse transcription (RT) was performed using the SuperScript™ IV CellsDirect™ cDNA Synthesis kit according to the supplier’s protocol (ThermoFisher Scientific). Relative RNA expression levels were quantified by quantitative PCR (qPCR) using the TaqMan™ Fast Universal PCR Master Mix (Applied Biosystems) and commercially available TaqMan™ probes for murine *Sod1* or human *Hprt* genes (Applied Biosystems). Expression data were analyzed using the ΔΔCq method normalized to the expression of duplex reference murine *RpL13* (Mm02526700_g1) and murine *RpL30* (Mm01611464_g1) genes for murine *Sod1* (Mm01344233_g1) and to the expression of duplex reference human *GAPDH* (Hs027558991_g1) and human *IPO8* (Hs0091457_m1) genes for human *HPRT* (Hs02800695_m1), based on raw quantification cycle (Cq) values and MIQE Guidelines (90), and are presented as mRNA levels relative to untreated cells. Experimental data (experimental triplicates, 6 to 8-point dose-response curves) are presented as means ± standard deviation (SD) and were analysed with GraphPad Prism^®^ software using a three-parameter log(inhibitor) vs. response nonlinear regression to determine the IC_50_.

### Care and handling of laboratory animals

Mouse studies – Procedures involving mice conform to National and European regulations (EU Directive N°2010/63) and to authorizations delivered to our animal facility (N° J 13 055 05) and to the APAFIS N°35462-2022021518324714-V2 and N° 33402-202110081456798-V4 by the French Ministry of Research and Local Ethics Committee (CE N°014). All efforts were made to reduce the number of animals and to minimize animal suffering. We used 8–14 week-old C57Bl/6 wild type (WT) male mice (Janvier-labs, Le Genest-Saint-Isle, France) and B-hTfR1 male mice (hTfR1-KI mice, Biocytogen, Beijing, China). Mice were acclimated for a minimum of 15 days before treatment administration.

Non-human primate (NHP) studies – Studies in olive baboons and cynomolgus monkeys were performed at the Station de Primatologie (SdP Rousset, UAR 846 - CNRS, France) and Cynbiose S.A.S. (Sainte-Consorce, France), respectively. Both test facilities were accredited by the Association for Assessment and Accreditation of Laboratory Animal Care (AAALAC). All experiments were conducted in accordance with the European Directive 2010/63/UE as published in the French Official Journal of February 7th, 2013. Naïve male olive baboons (*Papio anubis*, 2.60 – 4.50 kg), born and bred at the Station de Primatologie (SdP Rousset test facility, France) and aged 0.8-1.4 years old at the time of treatment initiation, were used for a preliminary exploratory study, under animal facility authorization N° C130877 and study authorization APAFIS N° 35743-202202101014381-V10 by the French Ministry of Research and Local Ethics Committee (CE N°071). Study animals were habituated to manipulations beforehand. They were separated from their usual social group only during treatment administration and biopsy collection (less than 2 hrs). Administration of test articles and muscle biopsies were performed under general anesthesia (ketamine 4 mg/kg and medetomidine 0.04 mg/kg given as an intramuscular injection) on a heating blanket. Following orotracheal intubation, animals were maintained under 1.5–3% sevoflurane with an oxygen flow rate of 1.5⍰L/min. Following treatment administration, animals were kept under observation during 48 hrs in a dedicated room accompanied by one of its social congeners, namely their mother whenever possible. Before muscle biopsies, analgesia was obtained by SC administration of meloxicam 0.2 mg/kg. Naïve male cynomolgus monkeys (Mauritian origin) aged 2.8 to 2.9 years old (2.85 – 5.55 kg) at the time of treatment initiation, were bred by Noveprim and transferred to Cynbiose S.A.S. test facility. The study was reviewed by the Animal Welfare Body of Cynbiose and the Ethics Committee of VetAgro-Sup (1 avenue Bourgelat, 69 280 Marcy l’Étoile, France) and approved under number 1465 (APAFIS N° 2016072117544328-V7). During the acclimation period and the *in vivo* experimental phase with cynomolgus monkeys, animals were housed in stainless-steel and plastic enclosure, as isosexual social groups (up to 6 animals of the same sex together), possibly with other congeners not included in the current study. Animals had an acclimation period of approximately 4 weeks, during which they were gradually trained to remain calm when being held by the operators during manipulations. This training program is aimed at mitigating the stress experienced by the animals during experimental procedures and covering rectal temperature measurement, blood sampling and intravenous dosing on restraint chair.

### Mouse and NHP studies and sampling procedures

WT or B-hTfR mice were dosed with vehicle (PBS) or test articles using either intravenous (IV) bolus at 5 mL/kg in the tail vein or subcutaneous (SC) bolus at 10 mL/kg between the shoulders. At prescribed time points, survival blood sampling was performed using the tail snip method, without exceeding a cumulative volume of 10% over one week and 20% over two weeks. For blood samples dedicated to plasma PK analysis, a volume of *ca*. 40-60 µL of blood were collected in Heparin-lithium coated tubes, immediately centrifuged at 2500×g for 10 min at 4°C and the plasma fraction was collected and stored at -80°C until analysis. At the final timepoint (terminal procedure), blood was withdrawn by a cardiac puncture method in mice under anesthesia with a mixture of ketamine (200 mg/kg) and xylazine (20 mg/kg) administered by IP route. Briefly, a hematocrit capillary (75mm/75µl, inner diameter 1.07mm - 1.23mm, Cat. 9100175, Hirschmann) was inserted into the right ventricle and a volume of *ca*. 80 µL of blood was collected in an EDTA-coated tube for haematology assessment (reticulocyte count). Then, a Lithium Heparin coated hematocrit capillary was used to collect a minimum of 100 µL of blood into dry tubes, and the plasma fraction was recovered as described above for PK analysis. Following blood sampling, mice were extensively perfused transcardially in the left ventricle with 60-80 mL of heparinized (10 UI/mL) NaCl 0.9% (RT) at a flow rate of 8-10 mL/min using a peristaltic pump, during 5-10min depending on quality of tissue discoloration. At the end of the procedure, organs were visually completely discoloured, with no trace of blood flowing out of the heart upon organs compression. At the end of the vascular wash procedure, the brain was removed from the skull, then transferred into a 35×10mm Petri dish (*e*.*g*. Falcon Ref. 353001) cooled on ice beforehand. Brain regionalization was performed on freshly collected brains to isolate the following regions from each hemibrain: parieto-temporal cortex, hippocampus, striatum, thalamus, hypothalamus, brainstem and cerebellum. Spinal cord was extracted according to a previously described procedure (91). Briefly, at the end of the vascular wash procedure, the entire spinal column is isolated from the rest of the animal and surrounding tissues. The spinal cord is isolated by hydraulic extrusion: a tip connected to a syringe filled with cold PBS is inserted into the distal end of the spinal column and the spinal cord is extruded by PBS flushing over a Petri dish cooled on ice beforehand. Isolated retinas were collected in a petri dish filled with PBS after removal of the anterior segment and the eye lens 0. 5 below the limbus and special attention was taken to remove as much as possible of the RPE (retinal pigment epithelium). For collection of bone marrow, femurs were carefully dissected after removal of adjacent skin and muscle tissues, dislocated at the hip, knee and ankle joints, and cleaned of any residual connective tissue. Following removal of the epiphyses, bones were placed into perforated 0.5 ml microcentrifuge tubes nested within 1.5 ml tubes and centrifuged at ≥ 10,000×g for 15 s to expel the bone marrow. For knockdown assessment using RT-qPCR, tissues, brain regions, isolated retinas and the cervico-thoracic part of the spinal cord were immediately immersed in Safe-Lock Eppendorf or cryopreservation tubes prefilled with at least 1 mL of NucleoProtect RNA preservation solution (Macherey-Nagel, Cat. 740400) and stored at -20 °C before further processing. For tissue PK analysis using Stem-Loop RT-qPCR, tissues, brain regions and the thoraco-lumbar part of the spinal cord were weighed and immediately frozen on dry ice and stored at -80 °C before further processing. For knockdown and distribution analysis on tissue sections using *in situ* hybridization (RNAScope® or miRNAScope®, respectively, ACD), hemibrains and lumbar part of the spinal cords were immediately frozen on dry ice and stored at -80 °C before further processing. After enucleation, freshly sampled eyes were immediately immersed in Hartman’s fixative solution (Sigma-Aldrich) for approximately 48h at RT. Following fixation, tissues were transferred to 70% ethanol for storage and held for up to four days until subsequent tissue processing (refer to histological analyses – FFPE tissue processing section).

Olive baboons received a 30-minute IV infusion of either PBS, an unconjugated siSOD1h-aminohexyl or a C5-siSOD1h conjugate generated as described hereabove. Twenty-eight and 84 days after treatment administration, approximately 100 mg punches (*ca*. 6 mm^3^, scalpel incision) were performed in the gastrocnemius, quadriceps and tibialis anterior muscles after exposing the muscle tissue (slight cutaneous scalpel incision). Following procedures, animals were returned to their social groups.

Cynomolgus monkeys, previously trained to undergo the following procedures in conscious state and seated in primate chairs, were dosed with PBS or the B8V32-hFc-siHPRT conjugate by 30-min IV infusion at 3 mL/kg through the saphenous veins (except one animal from the PBS group, which was dosed through the cephalic vein). At prescribed time points, blood was collected in conscious animals by puncture of a femoral vessel (venous and/or arterial blood), using a Vacutainer® kit. For plasma PK analysis and clinical chemistry, approximately 1 mL and 2 mL blood, respectively, were sampled into Lithium Heparin tubes and immediately homogenized by gentle manual inversion. For PK analysis and some Lithium Heparin tubes dedicated to clinical chemistry, plasma was extracted within 4 hrs after collection by centrifugation at 1800xg for 15 minutes at approximately +22°C, then stored at RT for clinical chemistry assessment within one day or stored at -80 °C until analysis. For haematology assessment, approximately 1 mL of blood was sampled into K3-EDTA tubes, immediately homogenized by gentle manual inversion and stored at RT pending analysis within 4 hrs after collection. At the final timepoint (terminal procedure), the animals were sedated by intramuscular injection of ketamine (5 to 10 mg/kg) and midazolam (0.5 mg/kg) and an analgesic (Buprenorphine, 0.03 mg/kg, IM) was administrated to reduce the possible pain caused by the exsanguination method. Euthanasia was performed by a lethal intravenous injection of a veterinary euthanizing agent. Immediately after ascertainment of death, a post-mortem laparotomy was performed, the right atrium was sectioned and complete exsanguination was conducted by manually flushing of ice cold NaCl or Ringer lactate containing heparin (1000 U/liter), via the heart’s left ventricle as far as possible up to the left atrium, for approximately 30 min. A peristaltic pomp was used at a flow rate of approximately 0.83 mL/s. The infusion continued until the fluid was clear and the organs appeared visually well flushed. Brains and spinal cords were extracted from the cranial cavity and vertebral column and placed in a chilled saline bath for approximately 3 minutes. Brains were sliced coronally (approx. 4 mm-thick slices) from the frontal part up to the cerebellar level using a brain cutting matrix for cynomolgus (Ted Pella, USA). Each slice was then bisected along the mid-sagittal plane into left and right half-slices. Spinal cords were cross-sectioned, and segments were collected at the cervical, thoracic and lumbar vertebral levels. For knockdown assessment using RT-qPCR, approximately 30 mg of peripheral tissues of interest and biopsy punches (approximately 3.5 mm diameter) taken from the right hemisphere of brain slices in specified regions were collected, immediately immersed in 2 mL cryotubes prefilled with at least 1 mL of NucleoProtect RNA preservation solution (Macherey-Nagel, Cat. 740400), kept on ice during the necropsy, then stored at approximately -80°C until analysis. For knock-down analysis on tissue sections using *in situ* hybridization (RNAScope®, ACD), left cerebral hemisphere half-slices were cut into one ventral and one dorsal parts and immediately immersed in 10% neutral buffer formalin (NBF) fixative solution for approximately 48h at RT. Following fixation, tissues were transferred to 70% ethanol for storage and held for up to four days until subsequent tissue processing (refer to histological analyses – FFPE tissue processing section).

### Target mRNA quantification in mouse and cynomolgus monkey tissues

After addition of a lysis reagent QIAZOL (QIAGEN), tissue samples (from 5mg, *e*.*g*. for retinas, up to *ca*. 200mg, *e*.*g*. for kidneys) were homogenized using a Precellys Evolution tissue homogenizer equipped with a Cryolys Evolution cooling system (Bertin Instruments). Chloroform was added to the homogenates, and they were centrifuged at 6000 rpm at 4 °C for 15 min. Aqueous phase was recovered, and total RNA was extracted using the RNeasy 96 QIAcube HT Kit (QIAGEN) in a QIAcube HT system (QIAGEN). RNA samples were analyzed and dosed using a Fragment Analyzer RNA kit (Agilent) in a Fragment Analyzer system (Agilent). Relative RNA expression levels were quantified by RT-qPCR in two steps, using at first, the High-Capacity RNA-to-cDNA kit (Applied Biosystems) for the RT and then the TaqMan™ Fast Universal PCR Master Mix (2X) no AmpErase™ UNG kit (Applied Biosystems) with commercially available TaqMan™ qPCR assays (Applied Biosystems) for murine *Sod1* (Mm01344233_g1), murine *Hprt* (Mm03024075-m1), murine *RpL13* (Mm02526700_g1), murine *RpL30* (Mm01611464_g1), Rhesus *HPRT* (Rh-02800695-m1), Macaca fascicularis *PPIB* (Mf02802985_m1), Macaca fascicularis *RPL13* (Mf-02807896-g1) and Macaca fascicularis *RPL30* (Mf02808057-g1) genes (Applied Biosystems) for the qPCR. The qPCR analysis was processed by CFX maestro software (BIORAD). Expression data were analyzed using the ΔΔCq method, based on raw quantification cycle (Cq) values and MIQE Guidelines (90), with normalization to the expression of *RpL13* and *RpL30* reference genes for mouse samples (multiplex qPCR), and to reference genes as indicated in Supplemental Table 1 for cynomolgus monkey samples (simplex qPCR). The results are expressed as means ± standard error of the mean (SEM) and are presented as mRNA levels relative to the control animals injected with PBS.

### Protein analysis using Western-Blot

For SOD1 protein analysis from mouse tissue samples, whole tissue lysates were prepared from mouse frozen tissues homogenized in PBS 0.25% Triton X-100 (Sigma, Cat. T8787) using a Precellys Evolution tissue homogenizer equipped with a Cryolys Evolution cooling system (Bertin Instruments). Lysis was completed by incubation on ice for 15-30 minutes in presence of a protease inhibitor cocktail (ref P8340, dil. 1/100e; Merck), followed by centrifugation at 13,000xg at 4°C for 10 minutes, and the supernatant was collected and stored at -80 °C until analysis. The sample protein concentration was measured using the DC™ protein assay kit (Cat. #5000111, Biorad). Five μg of total protein lysate was resolved by electrophoresis on Bis-Tris 4-12% polyacrylamide gels. One gel was prepared for Coomasssie staining to assess protein loading homogeneity and one gel was prepared for Western-Blot analysis. For Western-Blot, proteins were transferred onto a PVDF membrane using the iBlot3 transfer system (ThermoFisher Scientific). The membrane was blocked with TBS 0.1% Tween-20 (TBS-T) containing 5 % nonfat dry milk before incubation with anti-SOD1 antibody (rabbit anti-SOD1, Abcam, Cat. ab51254, 1:10,000 dilution in PBS-T-milk) overnight at 4°C. After washing with PBS-T, horeradish peroxidase (HRP)-conjugated donkey anti-rabbit secondary antibody (Jackson Immunoresearch, Cat. 711-035-152) was used (1:100,000 dilution in PBS-T-milk) for 1h at RT. Detection was performed using Trident femto Western HRP Substrate (Genetex, Cat. GTX14698) and GBox imaging (Genesys). Densitometric quantification was performed using either Image Lab 6.0.1 software (BioRad) or ImageJ.

Mouse and cynomolgus monkey TfR1 protein levels were evaluated from CNS regions and peripheral tissue samples isolated from WT mice and cynomolgus monkeys (PBS-injected monkeys from the study shown in Figure 7) as performed for SOD1 protein analysis. Following protein transfer onto a PVDF membrane, mouse and cynoTfR1 were revealed using a rabbit anti-TfR1 antibody (GeneTex, Cat. GTX102596), diluted 1/3,000 in Can Get Signal™ Immunoreaction Enhancer Solution 1 (Toyobo, Cat. NKB-101T), followed by an HRP-conjugated donkey anti-rabbit secondary antibody (Jackson Immunoresearch, Cat. 711-035-152), diluted 1:100,000 dilution in Can Get Signal™ Immunoreaction Enhancer Solution 2.

For determination of VHH-hFc-siRNA stability in rat tritosomes, samples from each incubation time point were diluted 1/200, to reach a theoretical 2 ng/µL concentration of the full construct, and 5 ng were loaded onto 4-12 % bis-tris polyacrymalide gels (ThermoFisher Scientific). Proteins were transferred onto a PVDF membrane using the iBlot3 transfer system (ThermoFisher Scientific). The membrane was blocked with TBS containing 0.1 % Tween-20 (TBS-T) and 5 % non-fat dry milk before incubation with antibodies overnight at 4 °C. The primary antibodies used here were HRP-conjugated anti-human Fc (Jackson Immunoresearch, Cat. 109-035-098, dil 1/10,0000) and an HRP-conjugated cocktail of rabbit monoclonal anti-VHH Abs (Genescript, Cat. A02026, dil 1/10,000). Primary antibodies were incubated for 1-2 hrs at RT. After washing with PBS-T, signals were detected using Trident femto Western HRP Substrate (Genetex, Cat. GTX14698) and Gbox imaging (Genesys).

### VHH-hFc quantification in mouse and cynomolgus monkey plasma samples using ELISA

#### Anti-hFc ELISA

For quantification of VHH-hFc-siRNA conjugates in plasma samples from WT mice, an anti-hFc sandwich ELISA method was used. Briefly, 96-well Maxisorp plates (ThermoFisher Scientific, Cat. 439454) were coated overnight at 4 °C with mouse anti-human Fc antibody (Jackson ImmunoResearch, Cat. 209-005-098) at 10 μg/mL in 0.1 M sodium bicarbonate buffer (Sigma, Cat. S8875-500G). After incubation, plates were washed five times with PBS containing 0.05 % Tween-20 (Euromedex, 2001-B). Wells were blocked with 150 μL of 1× casein (ThermoFisher Scientific, Cat. 37528) for 1 hr at RT with gentle agitation (600 rpm). After a further wash step, test samples and standards — prediluted as appropriate in PBS containing 0.1 % Triton X-100 (Sigma, Cat. T8787), 1/10 casein (ThermoFisher Scientific, Cat. 37528), a protease inhibitor cocktail (Sigma, Cat. P2714) and plasma matrix 1/200 — were added (50 μL/well) and incubated for 2 hrs at RT, 600 rpm. Plates were washed and subsequently incubated with 50 μL/well of HRP-conjugated anti-human Fc antibody (Jackson ImmunoResearch, Cat. 209-035-098) diluted 1:5000 in casein 1X for 1 hr at RT. After a final wash, 50 μL/well of TMB substrate (Sera-Care, Cat. 5120-0077) was added and incubated for 15 min at RT, protected from light. The reaction was stopped by adding 50 μL/well of 2 N H_2_SO_4_ (Sigma, Cat. 258105) and absorbance was read at 450 nm (Spectramax ID5, Softmax Pro 7.2). All samples and standards were analyzed at the minimum required dilution (MRD) for each matrix. Standard curves were fitted using a four-parameter logistic regression for quantification.

Anti-VHH-hFc ELISA. Quantification of VHH-hFc-siRNA conjugates in plasma samples from B-hTfR mice and NHPs was performed at Eurofins ADME BIOANALYSES using an anti-VHH capture followed by an anti-hFc detection. Briefly, 96-well Maxisorp plates (ThermoFisher Scientific, Cat. 439454) were coated overnight at 4 °C with MonoRab™ Rabbit Anti-Camelid VHH Cocktail (Genscript, Cat. A2014-200) at 1 μg/mL in 0.1 M sodium bicarbonate buffer. Plates were washed five times with PBS containing 0.05 % Tween-20 (Sigma, Cat. P7949), then blocked with 300 μL/well of 1X casein for 1 hr at RT with gentle agitation (300 rpm). After washing, test samples and standards — prediluted in PBS containing 1/10 casein (ThermoFisher Scientific, Cat. 37582), plasma matrix 1/50 — were added (100 μL/well) and incubated for 2 hrs at RT with gentle agitation (300 rpm). Plates were washed, then incubated with 100 μL/well of HRP-conjugated anti-human Fc antibody (Sigma-Aldrich, Cat. A0170) diluted at 0.2 µg/mL in casein 1X for 1 hr at RT. After a final wash, 100 μL/well of TMB substrate was added and incubated for 15 min at RT, protected from light. The reaction was stopped by adding 100 μL/well of 2 N H_2_SO_4_, and absorbance was read at 450 nm. All samples and standards were analyzed at the minimum required dilution for each matrix. Standard curves were fitted using a four-parameter logistic regression for quantification. Pharmacokinetic analysis of plasma profiles in mice and NHPs was performed using an independent model method (non-compartmental analysis) with the WinNonlin® software v8.4 (Pharsight Corporation, Pittsburgh, USA).

### siRNA quantification in mouse plasma and tissues using Stem-Loop RT-qPCR

Preparation of RNA extracts from plasma and tissue samples – Plasma samples were diluted in RNase free water (ThermoFischer scientific) using a minimum of 500-fold dilution to avoid matrix interference. Tissue samples were collected in 2 mL tubes containing ceramic beads (OZYME), immediately recovered by 1 ml of PBS (Phosphate Buffer Saline, ThermoFischer Scientific) containing 0.1 % Triton X-100 (Sigma, Cat. T8787) (PBS-T) and homogenized using a Precellys Evolution tissue homogenizer equipped with a Cryolys Evolution cooling system (Bertin Instruments). A volume of homogenates corresponding to 10 mg of tissue was recovered and placed into new 1.5 mL DNase/RNase free Eppendorf tubes and a sufficient volume of PBST was added to obtain a final concentration of 10 mg/mL of tissue. Diluted samples were incubated on a dry block at 95 °C for 10 min, vortexed, and placed on ice for 10 min before centrifugation at 16,000 g for 10 min at 4 °C. Supernatants were transferred to new 1.5 mL DNase/RNase-free Eppendorf tubes and analyzed immediately or frozen until analysis.

#### Preparation of calibration standards and quality controls – For mouse plasma

on the day of analysis, calibration standards were prepared from a 75 ng/mL siRNA solution, by performing five-fold serial dilutions in 500-fold diluted plasma in RNase free water (plasma recovered from control PBS-injected animals as blank matrix). Of note, calibration curves obtained using unconjugated siRNAs were perfectly superimposed to those obtained with VHH-hFc-siRNA conjugates, allowing unbiased quantification of the siRNA AS from both the intact conjugate and its release form. For mouse tissues: on the day of analysis, calibration standards were prepared with a 75 ng/mL siRNA solution, by performing five-fold serial dilutions in 100-fold diluted tissue lysates in RNase free water (tissue lysates recovered from control PBS-injected animals as blank matrix). Quality control samples (QCs) were prepared in plasma or tissue blank matrix at 2.5 to 2,500 pg/mL by performing ten-fold serial dilutions.

#### Stem-Loop reverse transcription

Twenty µL of samples (plasma or tissue lysates), calibration standards and QCs were transferred into a 96-well plate and placed into a pre-heated thermal cycler (SimpliAmp, ThermoFischer) at 95 °C for 10 min to denature the double-stranded siRNA and facilitate the annealing of the stem-loop RT primer (customized by ThermoFischer) to the antisense (AS) strand during the reverse transcription (RT) step. Each assay plate included at least 8 calibration standards (15 ng/mL to 192 fg/mL) and 4 QCs ranging from 2.5 to 2,500 pg/mL. To detect siRNAs using a TaqMan-based approach, we used a similar strategy as described in Chen et al. 2005 (92). Five µL of the denatured sample, calibration standard or QC was added to 10 µL of the RT reaction mixture (TaqMan™ MicroRNA Reverse transcription Kit-Applied Biosystems). Reactions were then performed by incubation at 16 °C for 30 min followed by 42 °C for 30 min, then stopped by inactivation at 85 °C for 5 min followed by incubation at 4 °C. Finally, a volume of 100.5 µL of RNase free water are added to the 15 µL of cDNAs obtained before running qPCR.

#### Quantitative PCR

Five µL of diluted cDNAs were added to 5 µL of qPCR Mix, containing TaqMan™ Fast Universal PCR Master Mix (2X) no AmpErase™ UNG kit (Applied Biosystems), and a custom TaqMan assay (probe and primers specific to the cDNA of interest, Applied Biosystems). The reaction mixture was then incubated at 50 °C for 2 min, followed by 95 °C for 20 s and 45 cycles at 95 °C / 2 s and 60 °C / 30 s. All reactions were duplicated. The qPCR analysis was processed using CFX maestro software (BIORAD) and the quantity of siRNA into each sample was back-calculated based on the equation obtained from the standard curve generated by CFX maestro software. Concentrations in ng/mL were then converted to nM using the corresponding siRNA molecular weight.

### Histological analyses by in situ hybridization (ISH)

#### Fresh frozen (FF) tissue processing

*FF* mouse brain and spinal cord samples were rapidly embedded in Optimal Cutting Temperature (OCT, CellPath) compound. Twenty μm sagittal cryosections of left-cerebral hemispheres or cross-sections of spinal cord were prepared using a cryostat at -20°C (Leica Biosystems). Slices were directly collected onto Superfrost Plus slides (Epredia) and then air-dried 30 min at RT, then for 1 h at -20 °C and stored at -80°C until staining.

#### Formalin-Fixed Paraffin-Embedded (FFPE) tissue processing

Mouse eyes were processed as described by Pang et al. (93) with minor modifications. Briefly, following fixation eyes were rinsed in PBS and, to facilitate fluid penetration and preserve morphology, two “windows” were notched into the eyeball: one in the anterior chamber and another in the posterior chamber behind the lens. These, along with 4 mm-thick fixed monkey brain slices, were dehydrated through a graded ethanol series (from 70% to 100%), cleared in xylene and paraffin-infiltrated using the HistoCore PEARL (Leica Biosystems) automated tissue processor in accordance with ACD guidelines for FFPE sample preparation. Tissues were embedded in paraffin wax using the Histocore Arcadia H station (Leica Biosystems), sectioned sagitally for the mouse eyes and coronally for the monkey brains at 4 µm on a microtome (Leica Biosystems) and mounted on Superfrost Plus slides. Slides were air-dried and then stored at RT prior to staining.

#### RNAScope and miRNAScope staining

Fluorescent RNAscope® assay was performed for mRNAs detection using the Multiplex Fluorescent Reagent Kit v.2 (ACD) and chromogenic miRNAscope® assay for siRNA detection using miRNAscope HD (RED) Assay Kit (ACD). Both assays followed the manufacturer’s protocols for FF and FFPE tissues. Briefly, FF sections were fixed in 10% NBF solution for 1.5 h at RT, dehydrated through a graded ethanol series (50% - 100%), incubated in H2O2 solution (ACD) for 10 min at RT and treated with Protease IV for 30 min at RT both assays. FFPE sections were baked (1 h at 60°C), deparaffinized in xylene and dehydrated in 100% ethanol. Following H2O2 treatment (10 min, RT), FFPE tissues underwent target retrieval (15 min at 99°C) and then digested for 30 min at 40°C with Protease Plus (ACD) for RNAscope or Protease III (ACD) for miRNAscope. All sections were hybridized for 2 h at 40°C with specific probes targeting mRNAs or siRNAs, alongside positive and negative controls to ensure RNA quality and signal specificity (**Supplemental Table 2)**. Following amplification steps, mRNAs were detected using Opal 570 or 650 fluorophores (Akoya Biosciences) and siRNAs with Fast Red detection reagent. Fluorescent slides were counterstained with DAPI (30 sec, RT, ACD) and mounted with ProLong Gold Antifade (Thermo Fisher Scientific). Chromogenic slides were counterstained with 50% Gill’s Hematoxylin (2 min, RT, Sigma-Aldrich), dried for 15 min at 60°C and mounted with EcoMount (BioCare Medical).

#### Microscopic analyses

Fluorescent and chromogenic large-scale images of brain and eye sections were acquired using a Zeiss Axio Scan.Z1 slide scanner equipped with a 20x objective (Plan-Apochromat 20x/0.8). Individual tiles were stitched automatically using Zen software (Zeiss) to generate whole-tissue overviews. Representative high-magnification insets were cropped from the resulting files for detailed visualization using Zen software (Zeiss). To accurately assess mRNA knockdown in specific neuronal populations, high-resolution confocal images were acquired using a Zeiss LSM 700 microscope. Images were taken with the 40x oil-immersion objective (EC Plan-Neofluar 40x/1.30). A minimum of two sections per tissue, per animal, and per condition were analyzed.

### Statistical analysis

All data were analyzed with GraphPad Prism v.10, using either a Student’s t test, one-way analysis of variance (ANOVA), or two-way ANOVA with multiple comparisons. Cellular elimination half-lives were estimated using a two-phase decay regression model and ED_50_ values of the dose-response relationships from *in vivo* studies were estimated using a three-parameters log vs. response nonlinear fit. Animal numbers involved in each shown *in vivo* study and any other statistical or graphical parameters are indicated in the corresponding figure legend. P-values < 0.05, < 0.01, < 0.001 and < 0.0001 were considered statistically significant and labeled with *, **, *** and ****, respectively, in the figures.

