## Supplemental Material for "Harnessing TfR1 for Cross-Species Systemic Delivery of siRNAs to Deep Brain Regions Using Single-Domain Antibodies"

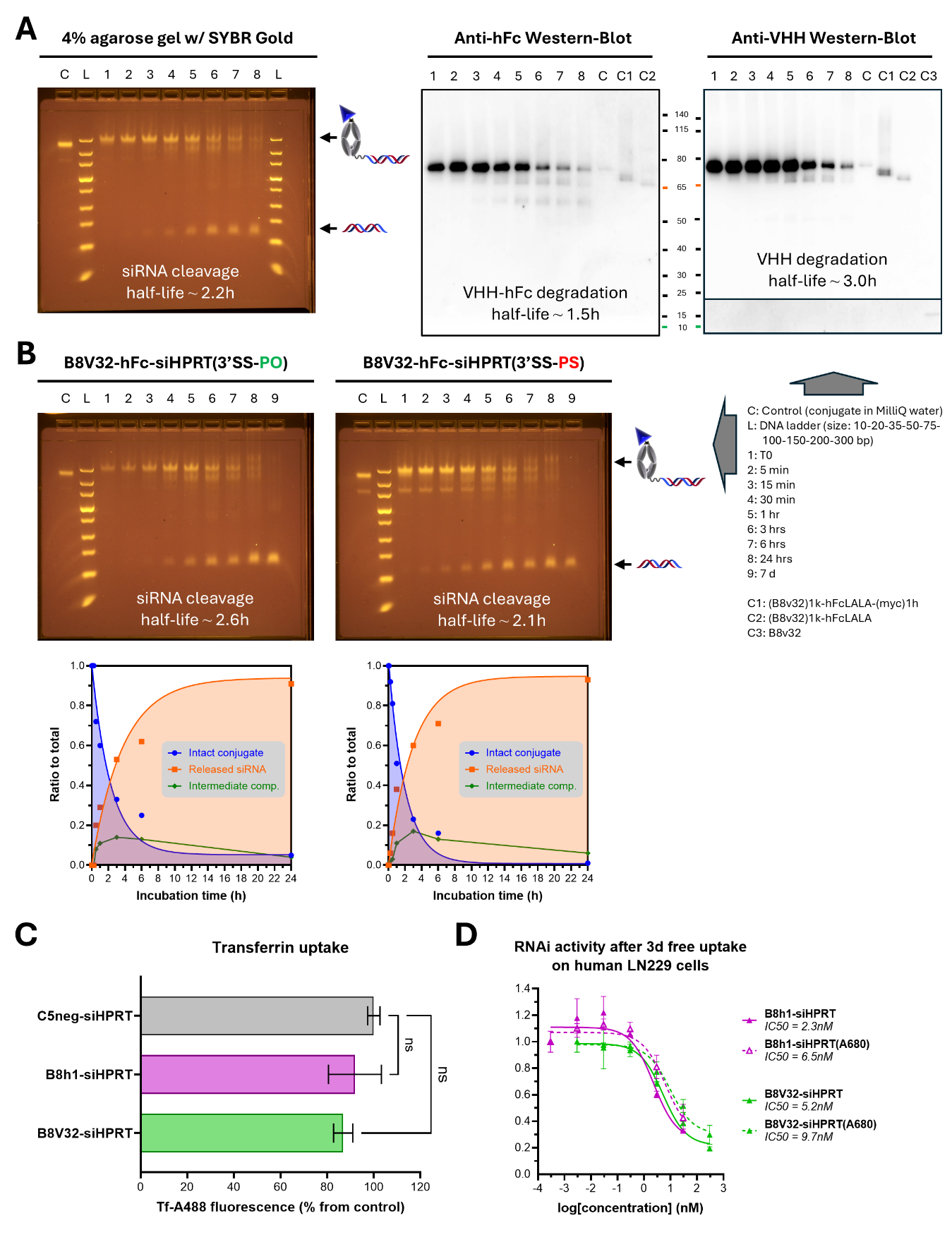

**Supplemental Figure 1. Intracellular metabolism and target engagement of A680-conjugated VHH-(hFc)-siRNA conjugates by free uptake *in vitro*.** (A) Time-course of the metabolic degradation of the B8V32-hFc-siHPRT conjugate in rat liver tritosomes, as assessed on both SYBR Gold pre-loaded 4% agarose gels (representative experiment shown in Figure 3E) and SDS-PAGE followed by anti-hFc or anti-VHH Western-Blot. (B) Comparison of the lysosomal metabolic profile of the B8V32-hFc-siHPRT conjugate comprising a PO- vs. PS-linker. Experimental data was generated from densitometry analysis of SYBR Gold pre-loaded 4% agarose gels; experimental curves showing the disappearance of the intact conjugate and appearance of the released siRNA were fitted using a nonlinear regression. (C) Transferrin uptake in human LN229 cells following pre-treatment with the indicated A680-conjugated VHH-siHPRT conjugates. (D) Target *Hprt* mRNA knockdown after 72 hrs free uptake of indicated unconjugated or A680-conjugated VHH-siHPRT conjugates on human LN229 cells. Results are expressed as means ± SD and experimental data were fitted using nonlinear regression.

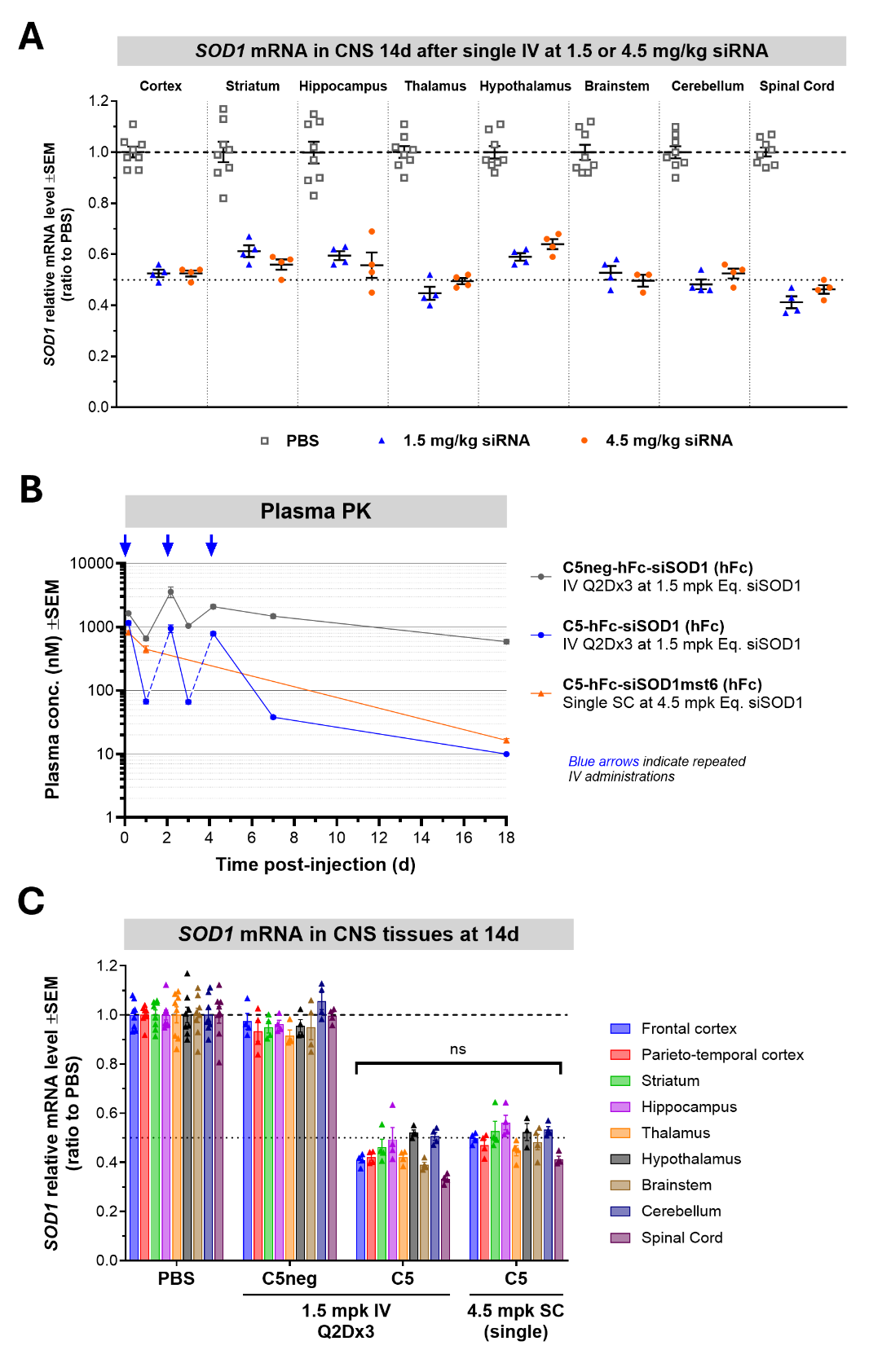

**Supplemental Figure 2. Multi-IV vs. single SC administration of a mTfR1-targeting C5-hFc-siSOD1 conjugate yields similar target engagement throughout CNS regions in WT mice.** (A) Shown is the *Sod1* mRNA KD in CNS regions 14 days after single IV administration of the C5-hFc-siSOD1 conjugate in WT mice at 1.5 or 4.5 mg/kg siRNA equivalent dose. (B) Plasma PK profile of the non-binding C5neg- or TfR1-binding C5-hFc-siSOD1 conjugate following repeated (Q2D x3) IV administration at 1.5 mg/kg siRNA or single SC administration at 4.5 mg/kg siRNA. *Sod1* mRNA KD in CNS regions 14 days after administration is shown in (C). Shown are means ± SEM.

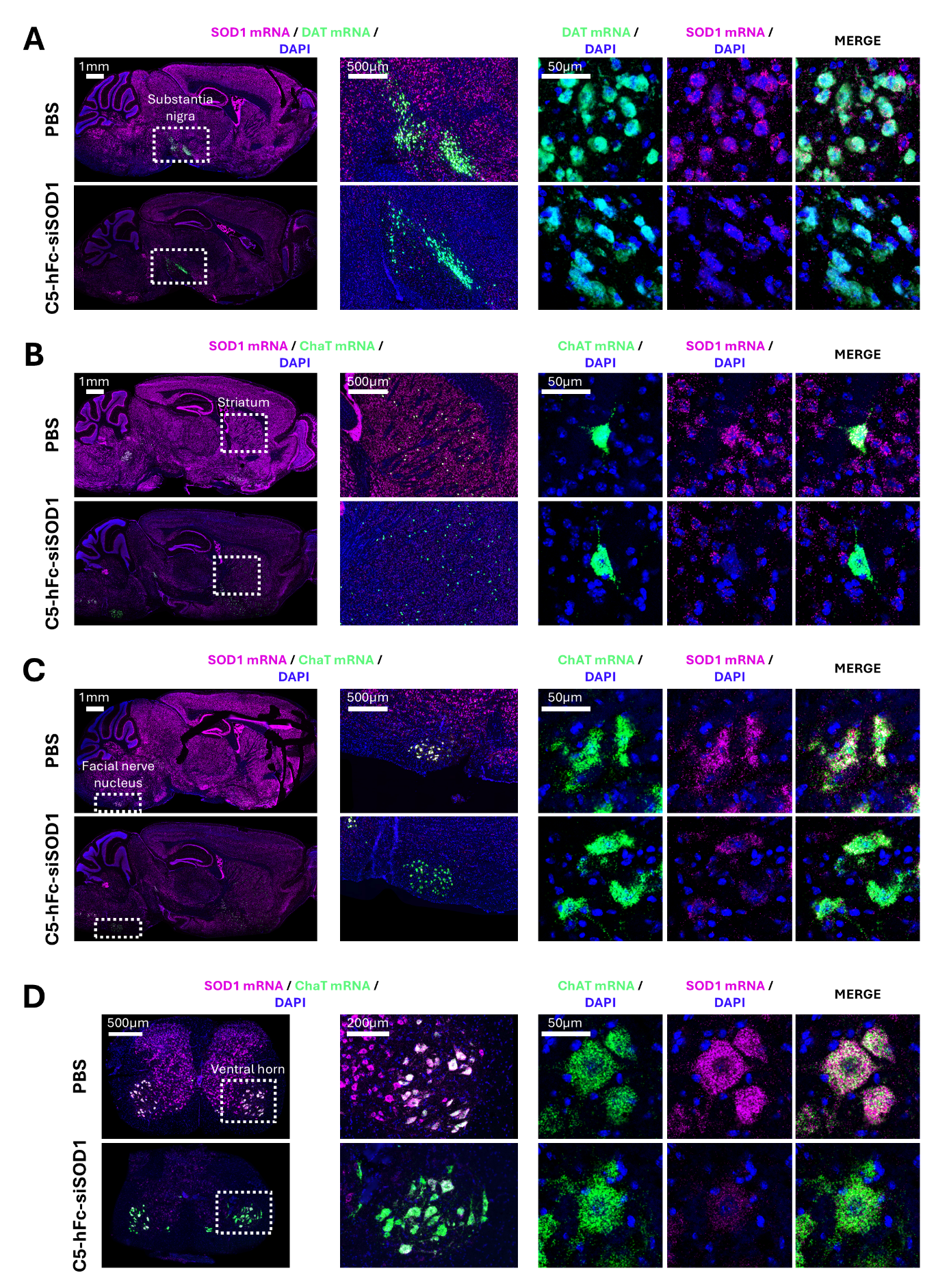

**Supplemental Figure 3. Systemic administration of a mTfR1-targeting C5-hFc-siSOD1 conjugate enables target knockdown across different neuronal populations throughout CNS regions in WT mice.** Shown are *in situ* hybridization micrographs and magnifications showing *Sod1* mRNA (RNAScope®, representative animal from n=4 mice per group) in dopaminergic neurons of the substantia nigra (A), cholinergic interneurons of the striatum (B), motor neurons of the facial nerve (cranial nerve VII) (C) and motor neurons in the ventral horn of the spinal cord grey matter (D), 14 days after multi-IV (Q2D x3) IV administration of PBS or the C5-hFc-siSOD1 conjugate at 1.5 mg/kg siRNA.

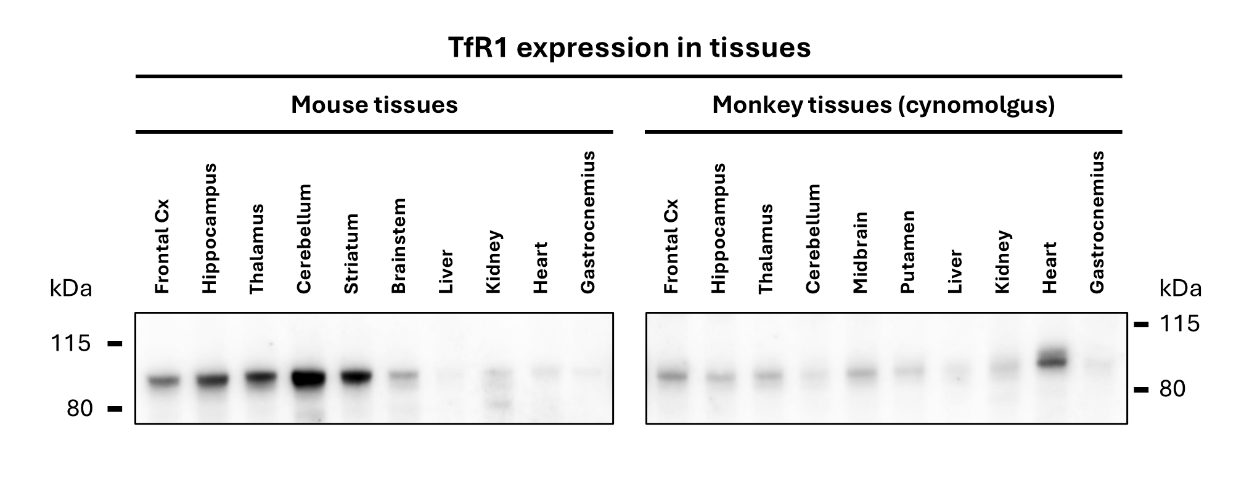

**Supplemental Figure 4. TfR1 protein expression across mouse and monkey tissues.** Shown are representative anti-mouseTfR1 or anti-cynoTfR1 Western-Blots, as indicated, from CNS regions and peripheral tissue samples isolated from WT mice and cynomolgus monkeys (PBS-injected monkeys from the study shown in Figure 7).

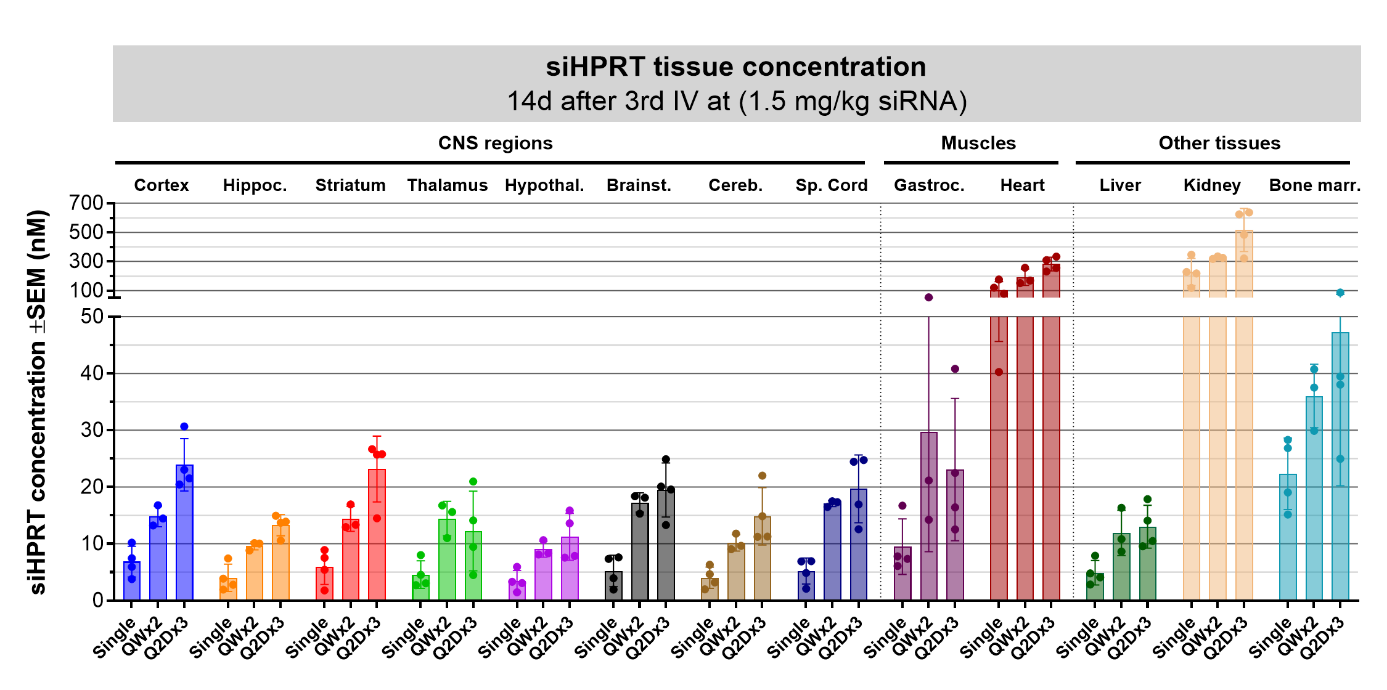

**Supplemental Figure 5. IV administration of a hTfR-targeting B8V32-hFc-siHPRT conjugate at low dose in hTfR1-KI mice achieves robust and dose-dependent accumulation throughout CNS regions.** Mice were dosed with the B8V32-hFc-siHPRT conjugate at 1.5 mg/kg siRNA equivalent dose as multi-IV bolus (Q2D x3). Shown are siRNA AS quantity in CNS and peripheral tissues, expressed as means ± SEM, 14 days after last IV dosing. *Hprt* mRNA levels are shown in Figure 5D.

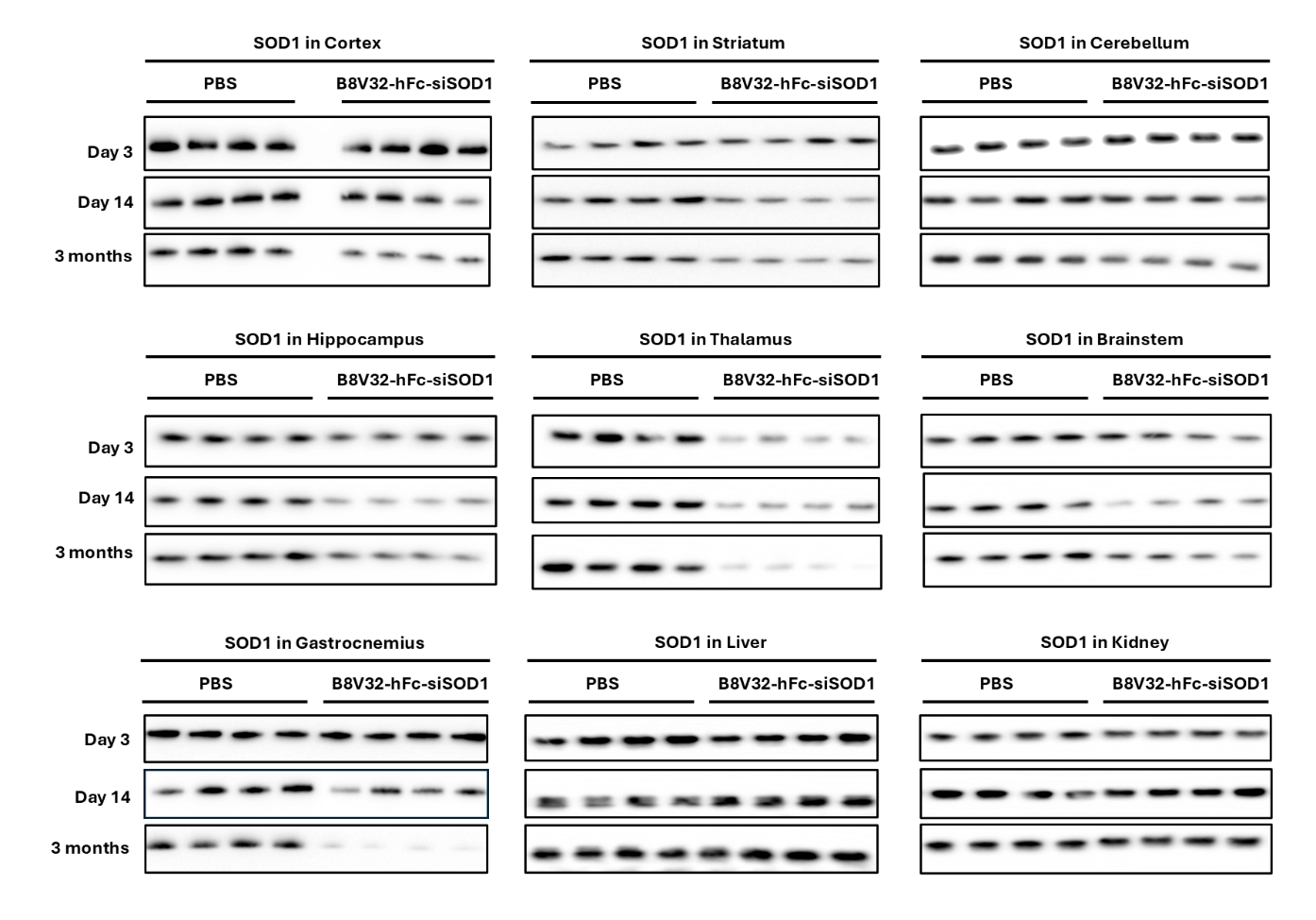

**Supplemental Figure 6. Time-course of SOD1 protein levels in CNS and peripheral tissues following treatment with the B8V32-hFc-siSOD1 conjugate in hTfR1-KI mice.** SOD1 protein levels were analyzed by immunoblotting of indicated tissue samples 3 days, 4 days or 90 days after multi-IV bolus (Q2D x3) of PBS or B8V32-hFc-siSOD1 at 1.5 mg/kg siRNA equivalent dose. Shown are individual results from 4 animals per group/timepoint.

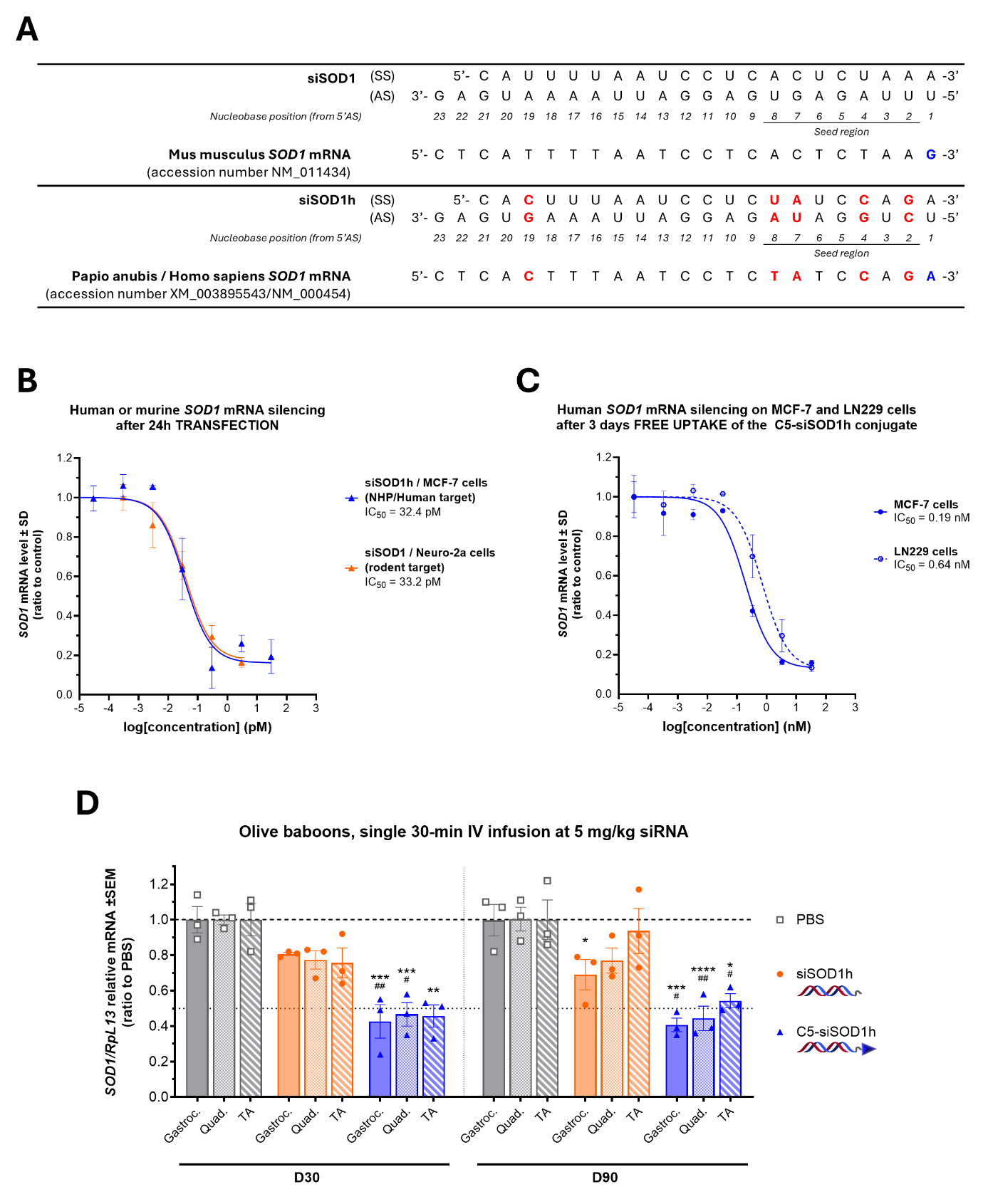

**Supplemental Figure 7. Time-course of the *SOD1* mRNA knockdown in muscle tissues after single IV administration of a C5-siSOD1h conjugate in olive baboons.** (A) The original siSOD1 targeting the rodent *Sod1* mRNA, as described by Brown and colleagues (5), was modified to allow full complementarity of the guide strand with the *SOD1* mRNA from olive baboons (*Papio anubis*). (B-C) Target *SOD1* mRNA knockdown 24h after transfection of the original siSOD1 on murine Neuro-2a cells or the engineered siSOD1h on human MCF-7 cells (B) or after 72h free uptake of the C5-siSOD1h conjugate on human MCF-7 or LN229 cells. Results are expressed as means ± SD and experimental data were fitted using nonlinear regression. (D) Olive baboons were dosed by single, 30-min IV infusion of the unconjugated siSOD1h or the TfR1-binding C5-siSOD1h conjugate. Muscle biopsies (gastrocnemius, quadriceps and tibialis anterior) were performed on each animal at one and three months after dosing, and *SOD1* mRNA levels were assessed using RT-qPCR. Results are expressed as means ± SEM.

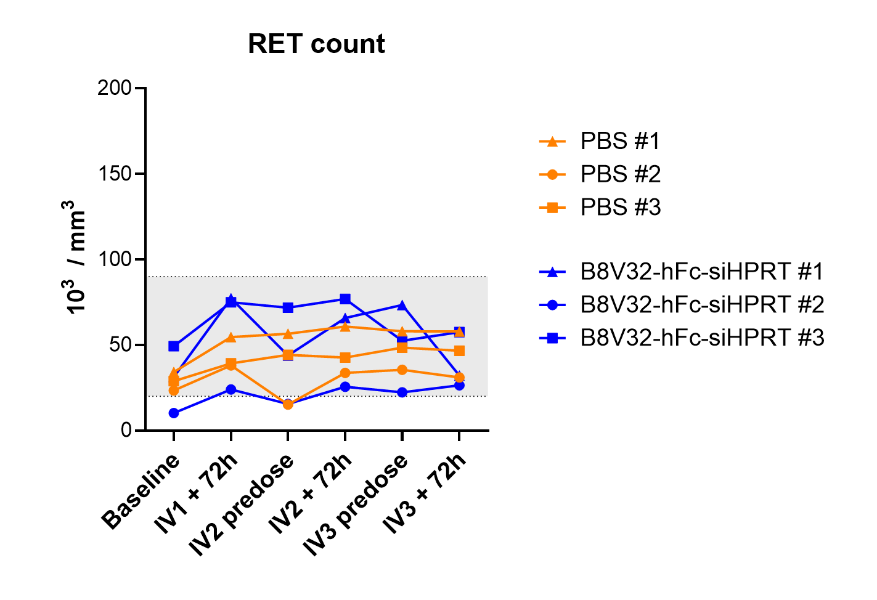

**Supplemental Figure 8. Reticulocyte count following repeated IV administration of the B8V32-hFc-siRNA conjugate in non-human primates.** Shown are individual reticulocyte count in blood samples from cynomolgus monkeys dosed by 30-min IV infusion three times at 7 days intervals with the vehicle (PBS) or with the B8V32-hFc-siHPRT conjugate (n=3 per group), at the same dose previously evaluated in hTfR-KI mice of 1.5 mg/kg siRNA equivalent dose.

**Supplementary Table 1.** List of reference genes used for normalization of the expression of the target HPRT mRNA in the NHP tissue samples.

| **Tissue sample** |  |  | **RpL30** |  | **RpL13** |  | **PPIB** |
| --- | --- | --- | --- | --- | --- | --- | --- |
| Frontal Cortex |  |  | X |  |  |  |  |
| Temporal Cortex |  |  | X |  |  |  |  |
| Caudate |  |  |  |  |  |  | X |
| Putamen |  |  |  |  |  |  | X |
| Globus Pallidus |  |  | X |  |  |  | X |
| Hippocampus |  |  | X |  |  |  |  |
| Thalamus |  |  | X |  |  |  |  |
| Hypothalamus |  |  | X |  |  |  | X |
| Midbrain |  |  | X |  |  |  |  |
| Pons |  |  |  |  |  |  | X |
| Medulla |  |  |  |  |  |  | X |
| Cerebellum Nuclei |  |  | X |  |  |  |  |
| Cervical Spinal Cord |  |  | X |  |  |  | X |
| Lumbar Spinal Cord |  |  | X |  |  |  | X |
| Liver |  |  | X |  |  |  | X |
| Kidney Cortex |  |  | X |  |  |  | X |
| Kidney Medulla |  |  | X |  |  |  | X |
| Lung |  |  |  |  |  |  | X |
| Spleen |  |  | X |  |  |  | X |
| Gastrocnemius |  |  | X |  | X |  |  |
| Diaphragm |  |  |  |  |  |  | X |
| Heart |  |  | X |  | X |  |  |

**Supplementary Table 2.** List of probes used for ISH experiments.

| **Platforms** |  | **Species** |  | **Targets** |  | **ACD catalog number** |
| --- | --- | --- | --- | --- | --- | --- |
| Manual Assay RNAscope |  | Macaca fascicularis |  | *HPRT1* mRNA |  | 584371 |
|  |  |  |  | *MAP2* mRNA |  | 1290331 |
|  |  |  |  | Positive Control - *POLR2A* mRNA |  | 500741 |
|  |  |  |  | Positive Control - *PPIB* mRNA |  | 424141 |
|  |  | Mus musculus |  | *SOD1* mRNA |  | 428581 |
|  |  |  |  | *MAP2* mRNA |  | 431151 |
|  |  |  |  | *CHAT* mRNA |  | 408731 |
|  |  |  |  | *DAT* mRNA |  | 315441 |
|  |  |  |  | *TTR* mRNA |  | 424171 |
|  |  |  |  | Positive Control - *POLR2A* mRNA |  | 312471 |
|  |  |  |  | Positive Control - *PPIB* mRNA |  | 313911 |
|  |  |  |  | Positive Control - *UBC* mRNA |  | 310771 |
|  |  | Bacillus subtilis |  | Negative Control – *DAPB* mRNA |  | 320871 |
| Manual Assay miRNAscope |  |  |  | siSOD1 antisense strand |  | 1142201 |
|  |  |  |  | siHPRT1 antisense strand |  | 1814141 |
|  |  |  |  | Negative Control - Scramble |  | 727881 |
|  |  |  |  | Positive Control - RNU6 |  | 727871 |
